# Polyclonal-Monoclonal Transition in Lung Squamous Cell Carcinoma Evolution

**DOI:** 10.64898/2026.03.27.714653

**Authors:** Taoyu Zhu, Yaping Xu, Junming Li, Zijin Wang, Zhenting Zhang, Boyu Wang, Mingyang Xiao, Boxin Liu, Manyu Xiao, Huiran Wang, Xinyi Xu, Ran Ji, Bo Yang, Songsong Li, Zuolin Shen, Xueqi Han, Xi Lu, Chen Lian, Xinyan Han, Yufei Liu, Silin Chen, Yunze Wang, Qian Tang, Yao Yao, Lian Wang, Huaqiong Huang, Qinglin Li, Da Wang, Xinwan Su, Bing Xia, Hongshan Guo, Xushen Xiong, Xuru Jin, Shirong Zhang, Yong Tang, Jian Liu

## Abstract

The evolutionary trajectory of lung squamous cell carcinoma (LUSC) remains poorly defined, hindering the development of effective therapies. By integrating genomic and transcriptomic sequencing from human LUSC specimens, we delineated a polyclonal-to-monoclonal evolutionary trajectory during LUSC progression. This evolutionary pattern was corroborated by single-cell RNA sequencing, which revealed consistent tumor cell heterogeneity. Specifically, the SBS5 mutational signature was enriched and correlated with poor prognosis independent of tumor stage. By further utilizing the spontaneous LUSC mouse model to identify key genomic and genetic events in LUSC progression, we observed that the JNK pathway was inhibited and that cytoskeleton-related pathways were dysregulated during LUSC development, and identified the mutations in the JNK pathway (e.g., *DACT1*) and cytoskeletal regulators (e.g., *KIF26A*). Collectively, these findings established a polyclonal–monoclonal evolution paradigm for LUSC, potentially regulated by JNK pathways, which could benefit LUSC precision therapeutics.

## Introduction

Lung cancer has the highest incidence and mortality among all cancers (Bray et al., 2022). It is categorized into small cell carcinoma, adenocarcinoma, squamous cell carcinoma, and large cell undifferentiated carcinoma (Gandara et al., 2015). Among these, lung squamous cell carcinoma (LUSC) accounts for 25% to 30% of lung cancer cases. To date, no targeted therapy has been developed explicitly for LUSC (Lau et al., 2022; Niu et al., 2022), while first-line immunochemotherapy (IC) demonstrates a modest efficacy, with a reported response rate of approximately 40% and unknown mechanisms of IC resistance (Zhou et al., 2021). In recent years, tumor evolution has been extensively studied in cancers such as hepatocellular carcinoma (Sun et al., 2024), colorectal cancer (Lu et al., 2024), and lung adenocarcinoma (Yang et al., 2022). However, similar research in LUSC remains lacking. Understanding LUSC evolution is critical for identifying key targets, thereby benefiting its therapy development.

Several tumor evolution patterns have been identified, including neutral evolution, branched evolution, linear evolution, convergent evolution, parallel evolution, and macroevolution (Vendramin et al., 2016; Yang et al., 2022). While these studies have characterized distinct tumor phylogenetics within individual patients, they have largely overlooked the broader evolutionary patterns occurring across different tumor progression stages. Consequently, the general evolutionary principles and their underlying driving mechanisms remain largely unexplored. Elucidating global evolutionary trends and their molecular mechanisms in LUSC could be instrumental in optimizing treatment timing and therapeutic strategies to improve patient outcomes.

A wide range of tumor mutational signatures has been reported, including single-base substitution signatures (Gelova et al., 2022), double-base substitution signatures, clustered-base substitution signatures, and insertion and deletion-related signatures (Alexandrov et al., 2020; Pei et al., 2020; Liu et al., 2024). While some mutational signatures have been well-characterized—such as SBS4, which is closely linked to tobacco smoking (Al et al., 2023), and SBS6, which is associated with defective DNA mismatch repair (Kim et al., 2023)—the clinical significance of many others remains poorly understood. Investigating the enrichment and prognostic relevance of mutational signatures in specific cancer types may enhance genomic subtype classification, refine prognostic prediction, and advance precision oncology.

Multiple LUSC subtypes have been proposed based on different classification criteria (Andre et al., 2024). Kadota et al. classified LUSC into keratinizing, non-keratinizing, and basaloid subtypes based on histological morphology (Kadota et al., 2014). Hammerman et al. categorized LUSC into four molecular subtypes: classic, primitive, basal, and secretory, using mutation and copy number variation data (Cancer Genome Atlas Research Network, 2012), later refined from a transcriptomic perspective (Wilkerson et al., 2010). Satpathy et al. applied proteomic analysis to classify LUSC into five subtypes: basal-inclusive, classical, EMT-enriched, inflamed-secretory, and mixed (Satpathy et al., 2021). Enfield et al. used tumor microenvironment profiling to define four distinct LUSC subtypes (Enfield et al., 2024). Despite these findings, no studies have integrated mutational signatures with distinct LUSC subtypes during LUSC evolution. Establishing these connections across multiple omics levels could provide a comprehensive understanding of LUSC biology, enabling precise classification and personalized therapeutic strategies.

By integrating multi-omics sequencing data from TCGA databases, TRACERx datasets, and other single-cell datasets, we identified a polyclonal–monoclonal evolution model for human LUSC progression. Specifically, we identified enrichment of the mutational signature SBS5, which correlated with poor prognosis in both transcriptomic and genomic analyses, and was also observed in the mouse LUSC model. Notably, we observed a distinct transition from secretory to basal and classic subtypes during LUSC progression, as revealed by transcriptomic profiling. Using our spontaneous LUSC mouse model to identify key genetic and genomic events during LUSC progression, we found that mutations in the JNK pathway (*e.g.*, *DACT1*) and alterations in cytoskeletal regulatory genes (*e.g.*, KIF26A downregulation) were pivotal to LUSC progression. We also observed an inhibited JNK pathway and dysregulated cytoskeleton-related pathways in LUSC, which might regulate the LUSC polyclonal-monoclonal transition.

## Methods

### Mouse models

*Pten*^f/f^*Lkb1*^f/f^ and CCSP^iCre^*Pten*^f/f^*Lkb1*^f/f,^ mice were utilized in this study. The CCSP^iCre^ mice were constructed following the procedures described previously (Vendramin et al., 2021) with some modifications. At the end of the administration, the mice were euthanized, and lung tissues were collected for the experiments or storage. All animal experiments followed the protocols approved by the Experimental Animal Center of the Zhejiang University-University of Edinburgh Institute, Zhejiang University.

### Samples collection and sequencing

Mice lung tissue samples were obtained from CCSP^iCre^*Pten*^f/f^*Lkb1*^f/f^ (experimental group) and *Pten*^f/f^*Lkb1*^f/f^ (control group) according to the stage of tumor development. We collected early tumor stage (ET, 2.5 months), small tumor (ST, 3.5 months), and big tumor (BT, 3.5 months) samples from CCSP^iCre^*Pten*^f/f^*Lkb1*^f/f^ mice. Corresponding control groups included early tumor stage control (EC, 2.5 months), and tumor stage control (TC, 3.5 months) samples, with n ≥ 6 biological replicates per group.

When the designated time points were reached, mice from each group were euthanized and lung tissues were collected. Mice lung tissue samples exceeding 120 mg wet weight were flash-frozen in liquid nitrogen in freezing tubes and stored at −80°C until sequencing preparation. All frozen samples were transported under dry ice conditions to ensure RNA integrity preservation during transit to the sequencing facility.

### RNA Extraction, library construction, and sequencing

Total RNA was extracted using Trizol reagent kit (Invitrogen, Carlsbad, CA, USA) according to the manufacturer’s protocol. RNA quality was checked using RNase free agarose gel electrophoresis. After total RNA was extracted, eukaryotic mRNA was enriched by Oligo(dT) beads. The enriched mRNA was fragmented into short fragments using the fragmentation buffer and reversely transcribed into cDNA by using NEBNext Ultra RNA Library Prep Kit for Illumina (NEB #7530, New England Biolabs, Ipswitch, MA, USA). Purified double-stranded cDNA fragments were end repaired, bases were added, and ligated to MGI sequencing adapters. Through the rolling circle amplification technology, a single DNA fragment is amplified into many copies to form a DNA nanoball (DNB). The ligation reaction was purified with the AMPure XP Beads (1.0X). Ligated fragments were subjected to size selection by agarose gel electrophoresis and PCR amplified. The cDNA library was sequenced utilizing DNBSEQ-T7 platform (MGI Tech) by Banian Medical Technology (Guangzhou) Co., Ltd.

### Histology staining

We performed H&E and immunohistochemistry (IHC) staining on mouse lung tissue sections to establish the landscape of LUSC progression. sections were deparaffinized, rehydrated, and then subjected to subsequent experiments. For H&E, the slides were stained with hematoxylin (Solarbio G1140, China) and eosin (Solarbio G1100, China). For IHC, the slides were conducted antigen retrieval at 100°C using a microwave for 10 minutes. Antigen retrieval was performed using Antigen Unmasking Solution (Vector Laboratories H-3300-250, USA) / 1 × TE Buffer. After cooling to room temperature, endogenous peroxidase activity was blocked using an IHC Kit (ZSGB PV-6000, China). 1% BSA was used for room temperature blocking for 1 hour. The primary antibodies and concentration used were anti-CK5 (Abcam ab52635, 1:1000, UK), anti-P40 (Biocare ACR3066A, 1:1000, USA), anti-TTF1 (DAKO M3575, 1:1000, UK), Finally, all slides were mounted using neutral balsam (Solarbio G8590, China).

### Single-cell sequencing

The preparation of single cells from mouse lung tissue was as follows: at the treatment endpoint, mice from each group were euthanized, and lung tissues were collected. After chopping with scissors, the tissues were digested using Tissue Dissociation Mix for 15 minutes. Once digestion was complete, the mixture was terminated and filtered through sterile filters (pore size: 40 μm). The resulting cell suspension was centrifuged at 350 g for 5 minutes to collect the pellet, which was then re-suspended and treated with red blood cell lysis buffer for 3 minutes. The final single-cell suspension was prepared, and lung single-cell mRNA was captured using the Singleron single-cell microfluidic workflow, ultimately generating cDNA for library construction. All procedures were supported by the Singleron single-cell platform. All dissociated cells were sequenced using the Singleron platform with 3’Reagent Kits following the manufacturer’s protocol. The single-cell library was then sequenced on the NovaSeq platform from Illumina PE150. CeleScope V1.14.1 was applied to process the barcodes and align the raw scRNA-seq data.

### Gene Set Enrichment Analysis (GSEA)

Gene Set Enrichment Analysis (GSEA) was performed to identify significantly enriched biological pathways and processes. The analysis was conducted using R (v.4.2.0) with the clusterProfiler package (v.4.10.1). A pre-ranked list of genes was generated based on the log2 fold change derived from our differential expression analysis. This gene list was then analyzed against the Hallmark and Canonical Pathways gene sets from the Molecular Signatures Database (MSigDB). The enrichment scores were computed with 1000 permutations.

### Alignment

Quality control was performed using FastQC (v.0.12.1). Fastp (v.0.24.0) was applied to remove adapters and low-quality reads. Trimmed reads were aligned to the GRCm38 genome assembly via BWA-MEM (v.0.7.18) (Alexandrov et al., 2020). Further quality control for BAM files was conducted via samtools (v.1.21) (Gelova et al., 2022).

### Somatic mutation calling

MuTect2 (Pei et al., 2020) was applied to call SNVs and indels, utilizing annotation files from GATK (v.4.2.0.0). Called variants were further filtered following the filtration parameter ‘PASS’. Function FilterMutectCalls and SelectVariants were applied for filtration. Vcf2maf was applied to annotate SNVs and indels following the Variant Effect Predictor (VEP100) database (Liu et al., 2024). Additionally, mutations were called using VarScan2, requiring a somatic P-value of ≤0.05. The sequencing depth in each region was also required to be at least 30, with a minimum of 10 sequence reads supporting the variant call.

### Removing mutations of low confidence

Since no paired tumor adjacent tissue was available, each sample was running for 6 times (each TC sample served as the control separately) during somatic mutation calling and copy number variation analysis. Specifically, for each tumor sample, it was compared to 6 biological replicate controls (e.g., ST1 with TC1-6) for SNVs and indels calling. The mutation was defined to be positive only if it passed the filtration after comparing to 6 control tissues separately, which was utilized for further analysis.

### Mutational signature extraction and deconvolution

Mutational signatures were extracted and deconvoluted as previously reported (Vendramin et al., 2021). The Hierarchical Dirichlet process (HDP) model was utilized via hdp package. The trinucleotide of the tumor was calculated and utilized as input data. Signatures that were previously reported to be enriched in lung cancer served as input, including SBS1, SBS2, SBS4, SBS5, SBS13, and SBS17b [55]. Hdp_init() function was utilized to initiate the model. Hdp_setdata() was used to assign trinucleotide to leaves, while dp_activate() was utilized to activate nodes. Hdp_posterior() was ran for 15 times to construct independent posterior chains. Hdp_multi_chain() was utilized to combined theses independent results, while hdp_extract_components() was utilized to extract components from them. Cosine similarity was calculated between components and reported signatures in the COSMIC database. For the robustness of the result, 5,000 iterations were utilized during de-nova signature extraction. The whole process was repeated twice to filter out false positive results.

NMF function was utilized to cluster TCGA LUSC patients according to mutational signatures. trincleotideMatrix() function was utilized to calculate the trinucleotide of the tumor. estimateSignatures() function was applied to decompose the matrix into several clusters while plotCophenetic() function was applied to determine the best cutoff, where k=5 was chosen here. After that, the signatures were extracted and calculated via extractSignatures() and compareSignatures() functions.

### LUSC selection pressure estimation utilizing dN/dS

Dndscv() function in the dNdscv package (Kim et al., 2023) was utilized to calculate the selection pressure from the cohort level. Default parameters were applied.

### Multi-hit mutations analysis

True positive multi-hit mutations were verified as previously reported (Vendramin et al., 2021). PyClone (Andre et al., 2024) was conducted to assign each mutation to clusters. After that, the evolutionary paths of each mutation were compared. If distinct mutations from the same gene did not overlap from the evolutionary aspect, they were regarded as true positive multi-hit mutations.

### Tumor phylogenetic trees construction and reconstruction

Pyclone was utilized for human phylogenetic trees construction, utilizing SNV data as input. For the analysis of humans, 166 samples were used (Stage I: 97, Stage II: 47, Stage III: 22), and the phylogenetic tree was constructed utilizing each sample within every patient. 3511 SNVs were included in the analysis by utilizing VarScan2 and Mutect mutation calls. After that, CONIPHER (Satpathy et al., 2021) was further applied to reconstruct phylogenetic trees for correction.

### Prognosis analysis

Kaplan-Meier Plotter (Gyorffy, 2024) (https://kmplot.com/analysis/) was applied to conduct the survival analysis, while the mean expression of selected genes was applied to compare the prognosis of patients with different levels of the SBS5-related transcriptomic signature. The Cox proportional hazards regression model indicated Significant differences by *P* values < 0.05.

### Pathway enrichment analysis

Both fgsea and clusterprofiler (Yu et al., 2012) were utilized for pathway enrichment analysis. Adjusted p <0.05 was regarded as the significantly enriched pathway. ggplot2 was utilized for visualization.

### scRNA-seq data quality control, processing, and annotation

Raw fastq data were firstly aligned to the mouse genome (GRCm38, ENSEMBL) by Celescope v1.14.1 following the CeleScope tutorial for scRNA-seq (https://github.com/singleron-RD/CeleScope/blob/master/doc/assay/multi_rna.md). Seurat (v4.1.1.9001) (Slovin et al., 2021) was applied to process the UMI count matrix. Doublets were assessed and removed by DoubletFinder (v2.0.3) (McGinnis et al., 2019). Both mitochondria and erythrocyte gene percentages were calculated by the PercentageFeatureSet function. The top 5000 genes were identified as highly variable genes (HVGs) using the FindVariableFeatures function of Seurat. All single cells were integrated according to sample ID by using SCTransform (Hafemeister and Satija, 2019). Dimensional reduction and clustering were conducted, where 1.5 resolution was selected for subcluster determination. Marker genes were determined using the FindAllMarker function of Seurat, while cell clusters were annotated as previously reported (Angelidis et al., 2019).

The dimension reduction and annotation of published human scRNA-seq data were downloaded from processed source data (Salcher et al., 2022). To ensure comparability across different time periods, we only retained human data that simultaneously contained Stage I, II, and III cases and originated from the same sequencing platform and batch.

The quality control related data of human published data and mouse data were listed in Table S3.

### Shannon ratio calculation

To quantify cellular heterogeneity during tumor progression, we calculated Shannon ratio as follows: After quality control and preprocessing of whole scRNA-seq data, tumor cells were extracted and processed by principal component analysis (PCA) for dimensionality reduction based on the top 2,000 highly variable genes. Unbiased clustering of all cells was subsequently carried out on these principal components. To compare the tumor cell heterogeneity, the Shannon diversity index was calculated based on the frequency distribution of tumor cells across the identified clusters, utilizing the diversity function of vegan (v2.6.8, with parameter ‘index=“shannon”’) (Oksanen et al., 2024). Finally, Shannon indices were compared across different stages to assess the dynamics of heterogeneity along tumor progression.

### RNA-seq pre-processing

Quality control was performed using FastQC (v.0.12.1) (https://github.com/s-andrews/FastQC). hisat2 (v.2.2.1) (Kim et al., 2019) was utilized to align the data to the GRCm38 genome. Samtools (v.1.21) (Li et al., 2009) and picard (v.3.0.0) (https://github.com/broadinstitute/picard) were utilized to deduplicate, sort, index, and normalize BAM files. featureCounts was utilized to count the reads. DESeq2 (Wilkerson et al., 2010) was utilized for differentially expressed genes calculation.

### Quantification and statistical analysis

*P* values < 0.05 were regarded as significant, and all statistical analyses were performed using R (v4.0.5, v4.2.2). Group comparisons were conducted via Wilcoxon rank-sum tests and ANOVA tests. Every experiment was repeated at least three times by using biologically independent samples.

## Results

### Polyclonal-to-Monoclonal Transition in Human Lung Squamous Cell Carcinoma (LUSC) Evolution

To depict the landscape of LUSC progression, we collected WES (Frankell et al., 2023), RNA-seq (CPTAC and TCGA), and single-cell RNA-seq (scRNA-seq) (Salcher et al., 2022) data from multiple datasets (*e.g.*, TCGA, CPTAC & TRACERX) (Fig.1A, Fig.S1A). Following quality control and stringent filtering, we observed that *TP53*, *CDKN2A*, *KMT2D*, *FAT1*, *PIK3CA,* and *NFE2L2* were the most frequently mutated genes in the cohort (Fig.S1B, Table S1), consistent with a previous publication [32]. We investigated the tumor evolution and selection pressure dynamics during LUSC progression. Via calculating the Simpson index for each phylogenetic tree generated from samples from each patient (Fig.1B-C, Fig.S2, S3A-B), revealing a significant decrease in phylogenetic complexity as LUSC advanced. We further validated our discoveries in human scRNA-seq data. After the quality control and preprocessing of published primary human LUSC single-cell data (Fig.S4A, Table S3), we extracted scRNA-seq data of primary tumor cells from stage I to stage III. To quantify the tumor cell heterogeneity, we calculated the Shannon distance based on their frequency distribution across clusters under unsupervised clustering (Fig.1D, detailed in Method). The decreased tumor cell heterogeneity during LUSC progression supported our findings (Fig.1D). To further quantify selection pressure, we calculated the ratio of non-synonymous to synonymous substitutions (dN/dS). The dN/dS ratio generally increased in Stage II and Stage III compared to Stage I for missense mutations (wmis) and all coding mutations (wall) (Fig.1E-F), and the dN/dS ratios of Stage II and Stage III are comparable. Because dN/dS was calculated at the cohort level, no statistical test could be conducted on it. This trend suggested progressively intensified positive selection during LUSC progression, particularly in the early phases of tumor evolution, hinting that strong positive selection potentially contributed to the monoclonal dominance in advanced-stage LUSC.

**Fig.1.**
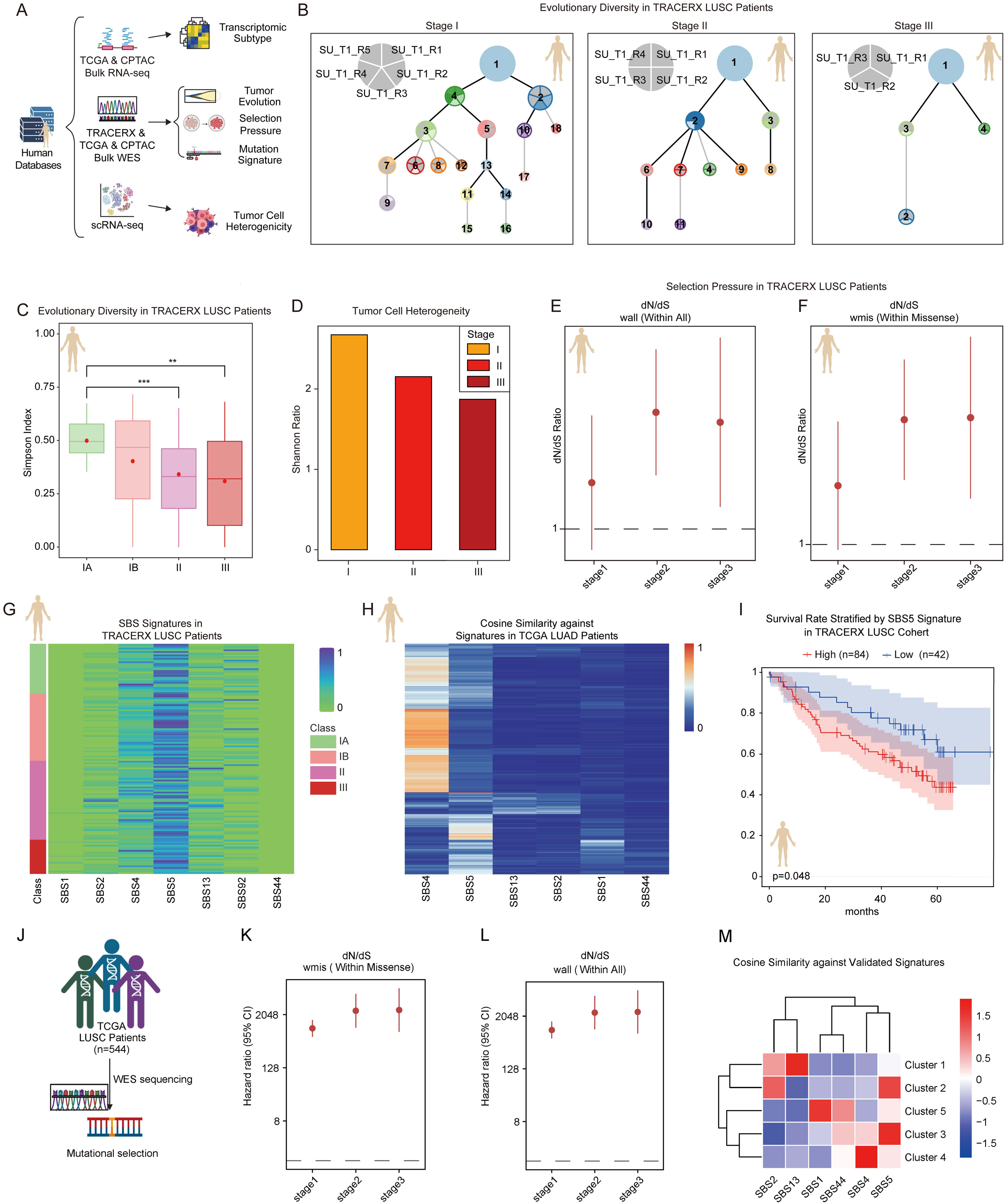
Polyclonal-to-Monoclonal Transition in Human Lung Squamous Cell Carcinoma (LUSC) Evolution. **A.** The workflow for collecting WES, bulk RNA-seq, and scRNA-seq data for transcriptomic and genomic analysis. **B.** Examples of representative phylogenetic trees in human Stage I, II III patients from TRACERX LUSC cohort. **C.** The box plot showing the Simpson index of phylogenetic trees across stages in TRACERX LUSC cohort. The dashed lines indicated the median values, while the red points indicated the mean values. Two-sided Wilcoxon test was applied for significance calculation. ** p < 0.01, *** p < 0.001. **D.** Bar plot showed the tumor Shannon distance across stages. **E-F.** Comparisons of missense point mutation and overall point mutation selection pressure across LUSC stages using dN/dS. **G.** Heatmap showed the score of reported NSCLC-related SBS signatures in patients from the TRACERX LUSC cohort across stages. **H.** Heatmap showed the score of reported NSCLC-related SBS signatures in patients from the TCGA LUAD cohort across stages. **I.** The Kaplan-Meier survival curves for overall survival (OS) in the TRACERX cohort LUSC and lung cancer patients. The x-axis represented the time in months, while the y-axis showed the probability of survival. **J.** The workflow for whole exon profiling of tumor samples from the TCGA LUSC cohort (n=544). **K-L.** Comparisons of missense point mutation and overall point mutation selection pressure across LUSC stages using dN/dS. **M.** Heatmap showed the score of reported NSCLC-related SBS signatures in patients from the TCGA LUSC cohort in distinct clusters.

To further characterize the mutational patterns in LUSC, we extracted the mutational signatures from TRACERX patients and compared them with reported signatures in LUSC patients (Gelova et al., 2022; Al et al., 2023). Among these signatures, SBS5 was significantly enriched in nearly all LUSC patients within the TRACERX cohort (Fig.1G). Initially, the SBS5 signature was identified as a clock-like signature (Alexandrov et al., 2015). Recent findings suggest that the SBS5 signature is particularly enriched in lung cancer patients exposed to high levels of PM2.5 (Hill et al., 2023). To verify whether SBS5 was also enriched in other cancer types, we conducted similar analyses in TCGA LUAD patients, in which we observed enrichment of the SBS4 signature, whereas the SBS5 signature was observed only in a subset of patients (Fig.1H). We further investigated the prognostic value of SBS5. Stratifying patients by SBS5 signature score revealed a significant association between SBS5 and poor prognosis (Fig.1I). Similarly, lung cancer patients with high SBS5 scores showed a trend toward poorer prognosis, although the difference was marginal (*p* = 0.054) (Fig.S4B). In summary, the SBS5 signature was enriched in human LUSC.

To confirm the robustness of our findings, we analyzed the whole-genome sequencing data from 544 LUSC patients in the TCGA database (Fig.1J). The dN/dS ratio exhibited an increasing trend for both wmis and wall mutations (Fig.1K-L), reinforcing our observation of progressively intensifying selection pressure during LUSC progression. Using non-negative matrix factorization (NMF) clustering, we stratified patients into five distinct clusters (Fig.S4C). Among these, Cluster 3 (n = 141 patients) exhibited significant enrichment of the SBS5 signature (Fig.1M), further supporting its prevalence in LUSC patients. These findings showed that a polyclonal-to-monoclonal transition in LUSC evolution occurred in more LUSC patients.

### Mouse LUSC Partially Mimics Human LUSC Evolution

To determine key factors in orienting the polyclonal-monoclonal transition in LUSC progression, we performed bulk whole-genome sequencing (WGS), bulk transcriptomic analysis, and scRNA-seq analyses on our spontaneous LUSC mouse model (Fig.2C), which we previously demonstrated to closely mimic human LUSC at both the transcriptomic and pathological levels [23] (Fig.2A-B). For example, morphological similarities between human and mouse LUSC were evident during LUSC progression (Fig.2A-B). In stage I of human LUSC, tumor tissue appeared in nests and strips, with tumor cells of varying sizes, observable nuclear division, and prominent nucleoli. Keratinization was evident, and scattered neutrophil infiltration was observed. In the early stage of mouse LUSC (ET), tumor tissue was arranged in nests, with the formation of keratin pearls. Tumor cells displayed variable sizes, occasional nucleoli, nuclear division, and scattered neutrophil infiltration (Fig.2A-B). In stage II of human LUSC, tumor tissue formed nests of polyhedral tumor cells, with numerous nuclear divisions, enlarged red nucleoli, and prominent intercellular bridges (Fig.2A-B). In a small tumor of mouse LUSC (ST), tumor tissue formed nest-like structures within the alveolar cavity, with tumor cells exhibiting irregular shapes, nuclear division, prominent nucleoli, and scattered neutrophil infiltration (Fig.2A-B). In stage III of human LUSC, tumor tissue formed sheet-like structures, with aneuploid tumor cells of variable size and less prominent nucleoli, and scattered lymphocyte infiltration (Fig.2A-B). In the big tumor of mouse LUSC (BT), tumor tissue appeared patchy, with polygonal tumor cells, visible nuclear division, and scattered neutrophil infiltration (Fig.2A-B). These findings highlighted the high histopathological similarity between human LUSC and our spontaneous mouse LUSC model during tumor progression, underscoring its reliability as a model for determining key factors in orienting the polyclonal-monoclonal transition in LUSC progression.

**Fig.2.**
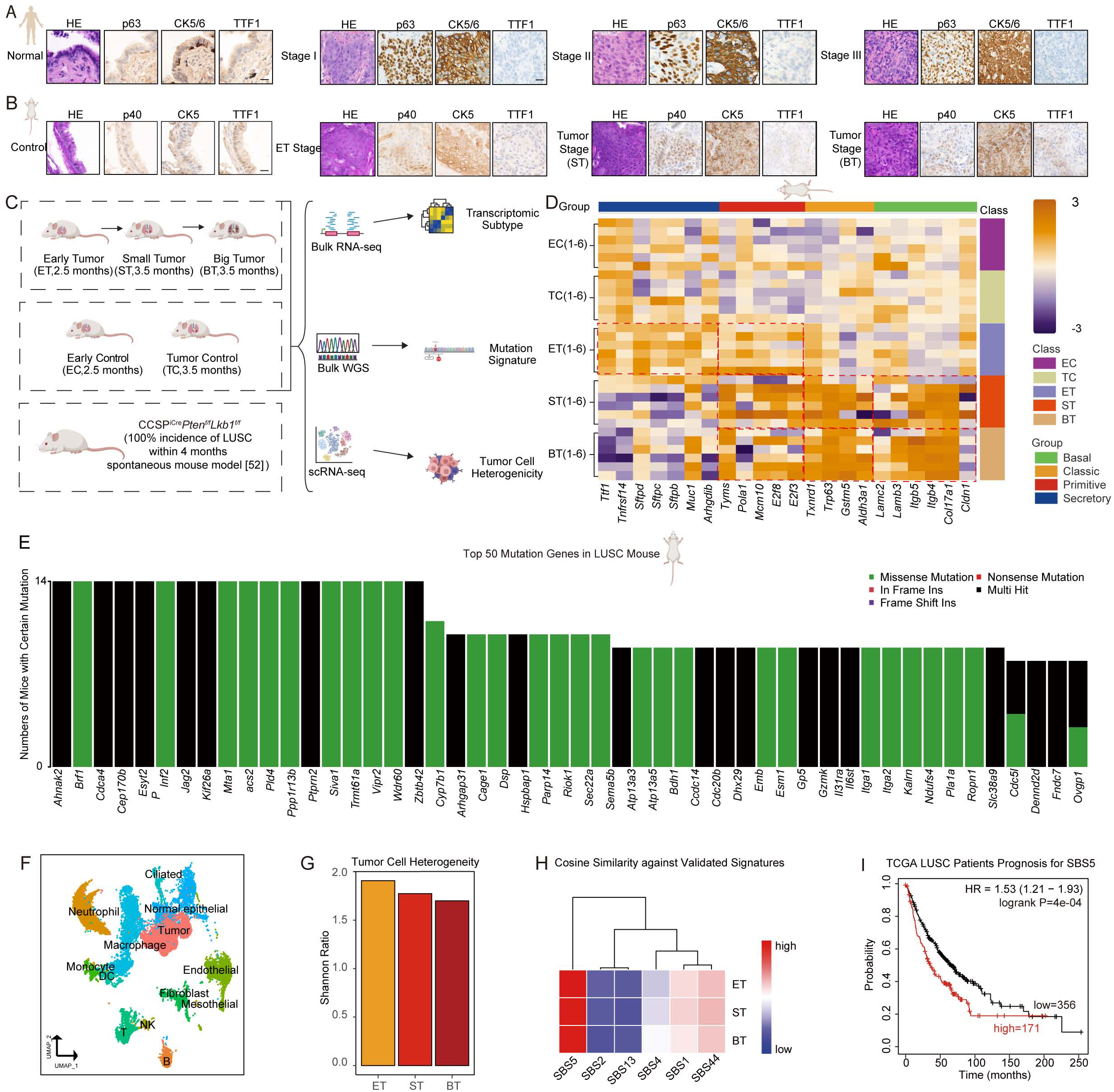
Mouse LUSC Partially Mimics Human LUSC Evolution. **A.** HE staining and IHC staining of P40, P63, CK5/CK6, and TTF1 in human LUSC samples. **B.** HE staining and IHC staining of P40, P63, CK5/CK6, and TTF1 in mouse LUSC samples. **C.** The workflow for whole genome and transcriptome profiling of mouse samples from ET (n=6), ST (n=6), and BT (n=6) stages. **D.** Heatmap showed the normalized expression level of reported LUSC transcriptomic subtype signature genes in the LUSC mouse model. **E.** Bar plot showed the top 50 mutated genes in mouse LUSC samples. **F.** UMAP showed annotated major cell types in mouse LUSC samples. **G.** Bar plot showed the tumor Shannon distance across stages. **H.** Heatmap showed the score of reported NSCLC-related SBS signatures in LUSC mouse model across stages. **I.** The Kaplan-Meier survival curves for overall survival (OS) in TRACERX cohort LUSC patients. The x-axis represented the time in months, while the y-axis showed the probability of survival. Scale bars, **(A)**;25μm **(B)**:20μm.

We further queried the transcriptomic similarity between human and mouse LUSC. Via comparing mouse transcriptomic profiling with reported transcriptomic subtypes in human LUSC (Wilkerson et al., 2010), we found that ET-stage tumors retained secretory and primitive subtype characteristics, which transitioned to primitive, basal, and classic subtypes in the ST and BT stages, with a predominant shift towards basal and classic subtypes (Fig.2D).

Following filtration and quality control, we identified abundant mutations in our spontaneous LUSC mouse model (Fig.2E, Table S2). Among them, *Ahnak2*, *Brf1*, and *Cdca4* were among the top three mutated genes. The *AHNAK2* mutation was reported to be a prognostic marker of immune checkpoint blockade efficacy in NSCLC (Cui et al., 2022). *BRF1* was reported to promote tumorigenesis in breast cancer and prostate cancer (Huang et al., 2019; Loveridge et al., 2020). *CDCA4* was also reported to be related to NSCLC progression (Xu et al., 2021; Shang et al., 2025). These results underscore the concordance between our spontaneous murine model and cancer patients, reinforcing the model’s relevance for further research.

To verify whether mouse LUSC mimics human LUSC evolution, we preprocessed our self-generated scRNA-seq data from mouse samples at ET and BT stages (Fig.2F, Table S3). We calculated the Shannon distance of tumor cells to quantify tumor heterogeneity similar to human LUSC in Fig.1D (detailed quality control and calculation in the Method section). We observed a decreased tumor cell heterogeneity during tumor progression (Fig.2G). These data, taken together, showed that mouse LUSC followed a polyclonal-monoclonal pattern (e.g., the decreased tumor cell heterogeneity) during progression.

We further extracted mutational signatures from our spontaneous mouse model as previously described (Al et al., 2023). We compared them with those reported in LUSC patients (Gelova et al., 2022; Al et al., 2023) to verify whether they showed similarities to human LUSC in terms of base mutations. Among these signatures, mutational signature analysis revealed a significant enrichment of the SBS5 signature across all stages (Fig.2H), indicating that our spontaneous LUSC mouse model mirrored a subset of LUSC patients. To further validate the clinical prognostic value of the SBS5 signature in LUSC patients, we derived a transcriptomic SBS5 signature based on the top dysregulated expressed genes (DEGs) in mouse LUSC progression and observed that patients with a high SBS5 transcriptomic signature had a significantly worse prognosis (Fig.2I). In summary, our spontaneous LUSC mouse model, previously demonstrated to closely mimic human LUSC at both the transcriptomic and pathological levels (Liu et al., 2019), progressed across multiple levels similar to human LUSC evolution, including the genome, transcriptome, and tumor cell heterogeneity, underscoring its value for further studying key genetic and genomic events during LUSC progression.

### JNK1/2 Pathway Was Mutated and Inhibited during LUSC Evolution

To identify key genomic events driving LUSC progression, we performed cross-species comparative analysis of human and mouse mutational landscapes across stages, identifying 53 conserved mutation genes (Fig.3A, Table 1). Pathway enrichment analysis in both human and mouse LUSC revealed that mutated genes were predominantly associated with immune regulatory pathways across all stages of progression. In the ET stage, pathways related to immune cell proliferation were significantly enriched, suggesting an active immune-tumor interaction. Enrichment in ion channel-related pathways characterized the ST stage. In contrast, the BT stage exhibited enriched LUSC development pathways, particularly the JNK pathway, which we previously identified as a key repressor of the LUSC progression, evidenced by *Jnk1/2* deletion significantly promoting the progression of mouse spontaneous LUSC tumors (Liu et al., 2019) (Fig.3B). We further identified key mutated genes associated with the downregulation of the JNK pathway during LUSC progression. Via overlapping genes enriched in JNK-related pathways in human LUSC samples and mouse BT samples, we identified three conserved genes, including *DACT1*, *NAIP*, and *ZMYND11* (Fig.3C). Among them, only *DACT1* was mutated in patients across all stages. Therefore, we conducted additional analyses of *DACT1* and observed nine mutations in human *DACT1* and one mutation in mouse *Dact1*, with distinct mutation rates (Fig.3D-E). Using AlphaFold3, we predicted the structural impact of *DACT1* mutations. We found that the mouse A562T mutation and human L104M, E221K, Q279H, P332R, N363Y, E429K, G517C, H586N, E688V, A721G, D800H, and K821Rfs*10 were predicted to have a significant impact on the overall protein structure of DACT1. In contrast, human S441L, Q592Rfs*100, and W784Lfs*30 were predicted to have negligible influence (Fig.3F-G, Fig.S5-6). We also quired the structural impact of mutations of NAIP and ZMYND11, whereas neglectable overall structural impact was observed for both human and mouse (Fig.3H-K, Fig.S7).

**Fig.3.**
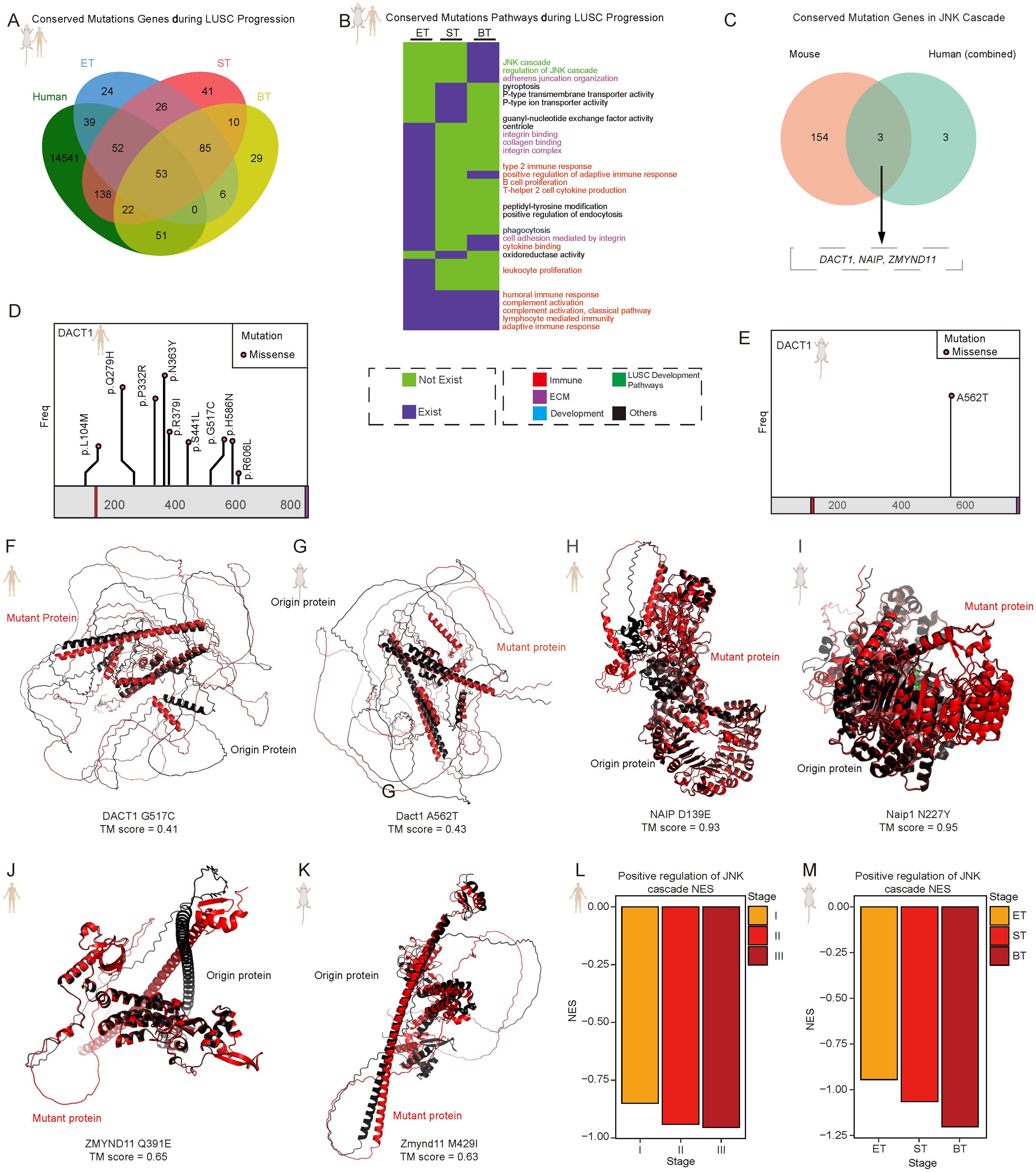
JNK1/2 Pathway Was Mutated and Inhibited during LUSC Evolution. **A.** The workflow of identifying conserved mutation genes during LUSC progression. **B.** Binary heatmap showed mutation genes conserved enriched pathways during LUSC progression. **C.** The Venn plot showed the workflow of identifying conserved mutation genes in JNK-related pathways. **D-E.** The Lollipop chart showed the position of mutations and variant allele fraction in the human and mouse DACT1 protein. **F-G.** Representative DACT1 whole protein structure variation in mouse and human predicted by AlphaFold3. TM score < 0.5 indicates significant variation. **H-I.** Representative NAIP whole protein structure variation in mouse and human predicted by AlphaFold3. TM score < 0.5 indicates significant variation. **J-K.** Representative ZMYND11 whole protein structure variation in mouse and human predicted by AlphaFold3. TM score < 0.5 indicates significant variation. **L-M.** Bar plot showed the NES of the positive regulation of the JNK pathway across stages in human and mouse samples.

To further verify our discovery, we conducted GSEA in both human and mouse single-cell analyses of tumor cells across stages, and we observed a decrease in JNK pathway activity during progression (Fig.3L-M). This was consistent with our published results in human and mouse LUSC tumors (Liu et al., 2019). These findings altogether showed that the JNK pathway was inhibited during LUSC progression, while *DACT1* mutation might mediate this process.

### Dysregulation of Cytoskeleton-Related Pathways in LUSC Evolution

To identify potentially key transcriptomic processes driving LUSC progression, we performed cross-species DEG analysis in both human and mouse LUSC across stages, revealing 2,079 conserved DEGs (Fig.4A, Table 3). Pathway enrichment analysis of conserved DEGs revealed significant enrichment in immune regulation and morphogenesis-related pathways across all stages of LUSC progression. In the ET stage, key pathways, including immune regulation, metabolic reprogramming, cytoskeletal dynamics, and classical oncogenic signaling (*e.g.*, cell cycle regulation and p53), were significantly enriched. In the BT stage, protein synthesis-related pathways were significantly enriched, indicating elevated protein biosynthesis activity, which may contribute to tumor progression and growth (Fig.4B).

**Fig.4.**
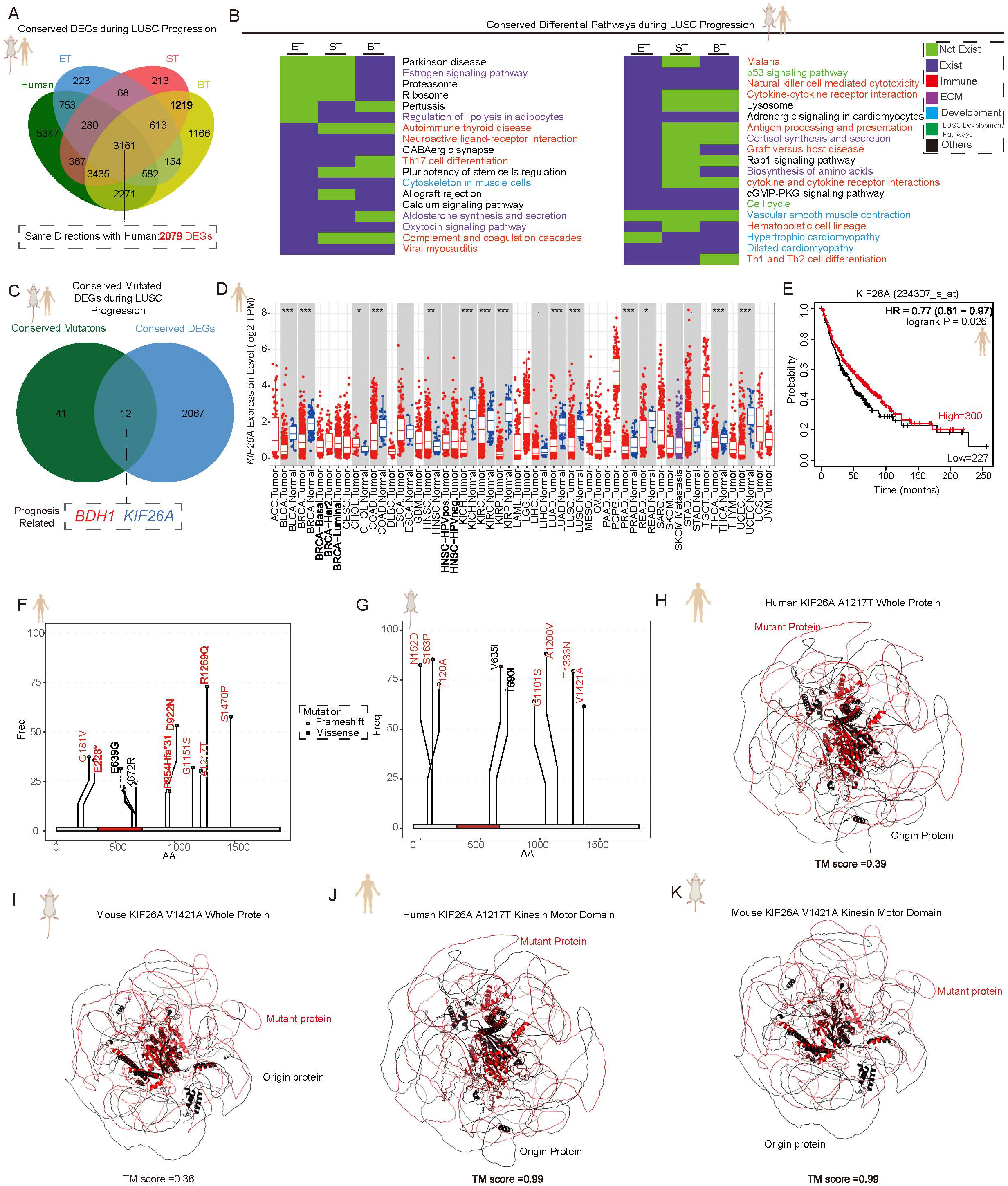
Dysregulation of Cytoskeleton-Related Pathways in LUSC Evolution. **A.** The workflow of identifying conserved differentially expressed genes during LUSC progression. **B.** Binary heatmap showed mutation genes conserved enriched pathways during LUSC progression. **C.** The workflow of identifying conserved differentially expressed mutated genes during LUSC progression. **D.** The box plot showing the expression level of KIF26A in the TCGA database in pan-cancer. The dashed lines indicated the median values. A two-sided Wilcoxon test was applied for significance calculation. * p < 0.05, ** p < 0.01, *** p < 0.001. **E.** The Kaplan-Meier survival curves for overall survival (OS) in the TCGA cohort of LUSC patients. The x-axis represented the time in months, while the y-axis showed the probability of survival. **F-G.** The Lollipop chart showed the position of mutations and variant allele fraction in the human and mouse KIF26A protein. **H-I.** Representative KIF26A whole protein structure variation in mouse and human predicted by AlphaFold3. **J-K.** KIF26A motor domain structure variation in mouse and human predicted by AlphaFold3. TM score < 0.5 indicates significant variation.

We further investigated conserved DEGs regulated by mutations. Through overlap analysis, we identified 12 candidate genes, whereas we found that *BDH1* and *KIF26A* were significantly correlated with prognosis in LUSC patients (Fig.4C, 4E, S8B). Among these, *KIF26A* was significantly downregulated (Fig.4D), and its low expression correlated with poor prognosis (Fig.4E). Although *BDH1* was significantly upregulated in LUSC samples (Fig.S8A), its high expression correlated with a favorable prognosis (Fig.S8B). Therefore, *KIF26A* was selected for further analysis. KIF26A harbored 10 mutations in human samples and nine mutations in mouse models (Fig.4F-G). Interestingly, major structural alterations were observed in KIF26A following mutation at the whole-protein level (Fig.4H-I, Fig.S8C, S9A-C, Table 2). In contrast, only negligible changes were detected in the motor domain region (Fig.4J-K, Fig.S9D-E, S10, Table 2). Further structural analysis revealed that KIF26A contained a high proportion of disordered regions, which may facilitate phase separation. Based on multiple predictive models, KIF26A was highly likely to undergo phase separation (Table 4). Therefore, we hypothesized that *KIF26A* mutations might contribute to LUSC initiation and progression via a phase-separation-dependent mechanism.

In summary, the progression of LUSC followed an ‘increase’ pattern of selection pressure, leading to a corresponding polyclonal-to-monoclonal transition in tumor clonal diversity, as reflected in tumor cell heterogeneity. The SBS5 mutational signature was highly enriched in LUSC patients and strongly correlated with poor prognosis. At the transcriptomic level, LUSC exhibited a transition from the secretory subtype to the basal and classical subtypes during initiation and progression. At the pathway level, the JNK signaling and cytoskeleton pathways likely play key roles in LUSC progression. Additionally, cytokine-cell interaction pathways were prominent in the early tumor stage. As LUSC progressed, metabolic reprogramming, ion channel activity, and classical oncogenic pathways became increasingly enriched (Fig.5).

**Fig.5.**
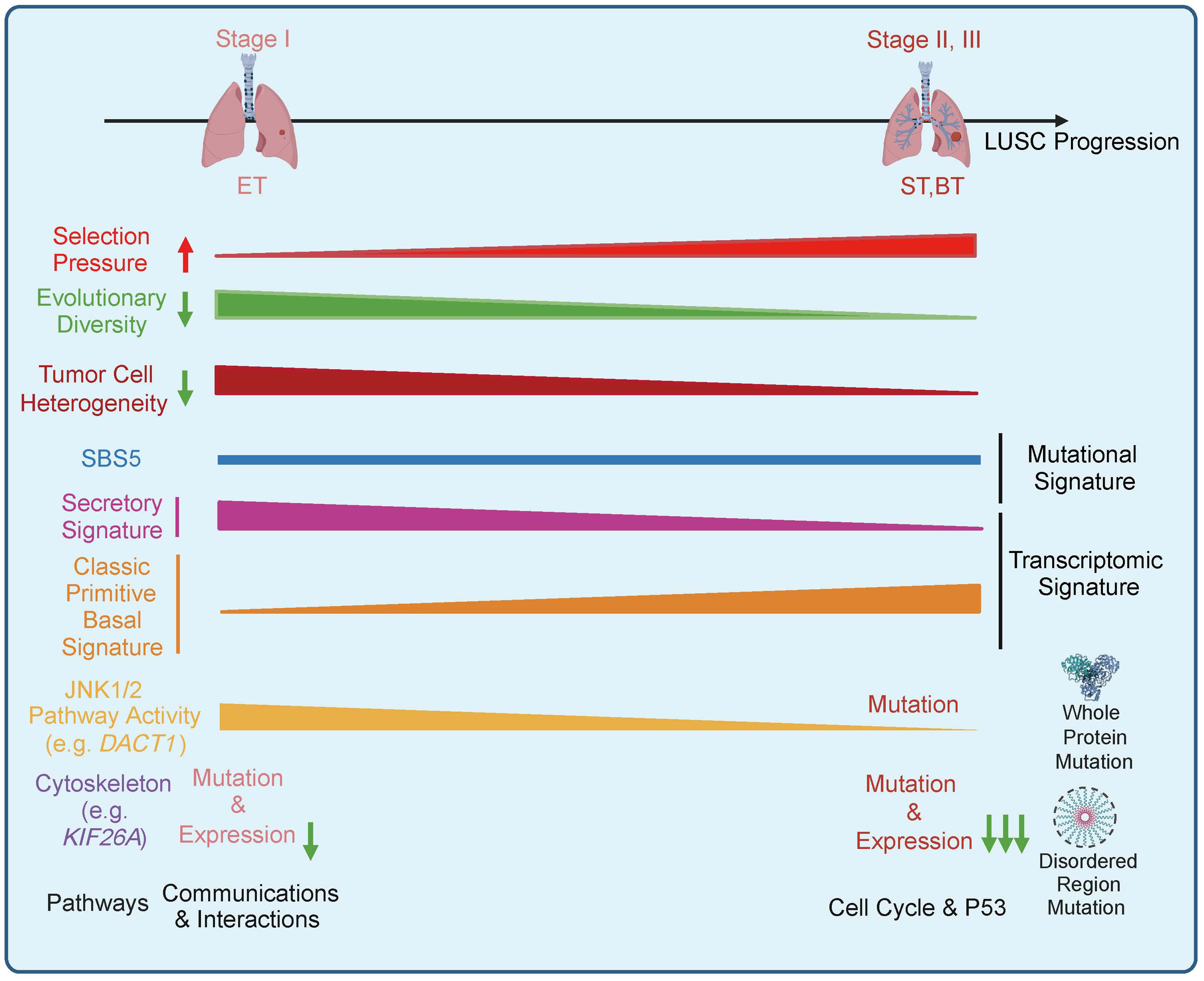
Working model. The progression of LUSC followed an ‘increase’ pattern in selection pressure, leading to a corresponding ‘decrease’ trend in tumor clonal diversity, which is supported by the single-cell transcriptomic analysis. The SBS5 mutational signature was highly enriched in LUSC patients and showed a strong correlation with poor prognosis. At the transcriptomic level, LUSC exhibited a transition from the secretory subtype to the basal and classical subtypes during initiation and progression. At the pathway level, the JNK signaling and cytoskeleton-related pathways likely play key roles in the progression of LUSC. Additionally, cytokine-cell interaction pathways were prominent in the early tumor stage. As LUSC progressed, metabolic reprogramming, ion channel activity, and classical oncogenic pathways became increasingly enriched.

## Discussion

By integrating genomic and transcriptomic analyses, this study delineated the key evolutionary trajectories and driving mechanisms underlying LUSC progression. We identified a clear evolutionary pattern characterized by a transition from polyclonal origins towards monoclonal dominance, accompanied by enrichment of a distinct SBS5 mutational signature and a phenotypic shift from the secretory subtype toward basal/classical subtypes. Early-stage disease was marked by widespread metabolic reprogramming and activation of immune-related pathways. In contrast, late-stage progression was closely associated with the accumulation of mutations in the JNK signaling pathway (*e.g.*, *DACT1*) and cytoskeletal regulators (*e.g.*, *KIF26A*). Collectively, these findings bridged the macroscopic clinical behavior of LUSC with the microscopic cellular and molecular evolutionary dynamics driving tumor progression.

Here, we demonstrated that LUSC followed a polyclonal–monoclonal evolution model, supported by phylogenetic tree construction and Simpson index analysis to assess tumor complexity. Our results further suggested that selection pressure dynamics might drive these evolutionary shifts. Interestingly, our findings aligned with the evolutionary patterns observed in colorectal cancer (Lu et al., 2024). Specifically, both our paper and theirs have found that tumors showed a decreased clonal diversity during progression. In colorectal cancer, multiple independently originated clones coexist and compete during the benign stage, with eventual outgrowth of a single dominant clone that evolves into carcinoma (Lu et al., 2024). The observation of a similar evolutionary paradigm in LUSC suggests that such a pattern may represent a standard early evolutionary route in epithelial cancers with complex tissue architecture. This evolutionary pattern demonstrates that early tumor heterogeneity and clonal competition are the basis for subsequent malignant progression.

Moreover, Lu et al. reported that colorectal cancer cells exhibited strong interactions with cells within the TME during the inflammatory stage (Lu et al., 2024). Similarly, we also observed significant enrichment of cell-cell interaction-related pathways in the ET stage. Therefore, such heterogeneity might be attributed to spatial heterogeneity during LUSC progression, in which cancer cells interact with distinct cell states within the tumor microenvironment. Distinct selection pressures imposed by both extrinsic factors, such as cell-cell communication, and intrinsic factors, such as tobacco-induced mutations, could both contribute to the emergence of polyclonal populations (Li et al., 2020). However, before distal or metastatic dissemination, a stringent bottleneck that requires the simultaneous acquisition of invasion and immune evasion capacity restricts dominance to a single subclone, thereby restoring monoclonality (Muller et al., 2016; Picard et al., 2020; Gharib et al., 2024), although further research is warranted to verify this hypothesis.

The polyclonal–monoclonal trajectory we observed in LUSC, mirroring dominant-clone dynamics, was also reported across multiple solid tumors and underscored the nonlinear evolutionary course of tumors. For example, Van Egeren et al. showed that high proportions of benign and dysplastic premalignant polyps were polyclonal in origin among 123 samples from six individuals with familial adenomatous polyposis (Van et al., 2025). Gao et al. demonstrated that most copy-number alterations in triple-negative breast cancer were acquired early, followed by polyclonal expansion during progression and final clonal dominance at metastasis (Gao et al., 2016). Furthermore, similar polyclonal–monoclonal evolutionary patterns have also been described in ovarian cancer, prostate cancer, and clear-cell renal-cell carcinoma patients (Venkatesan et al., 2016; Williams et al., 2016; Williams et al., 2018), extending the universality of our observations beyond LUSC.

A wide array of tumor mutational signatures has been identified (Gandara et al., 2015; Gelova et al., 2022). However, their correlation with transcriptomic alterations and prognostic relevance remains unclear. We identified a significant enrichment of the SBS5 mutational signature in a substantial proportion of LUSC patients. By calculating the *de novo* SBS5 signature enrichment score, we demonstrated its prognostic significance in both LUSC and broader lung cancer cohorts. Additionally, using our paired multi-omics spontaneous LUSC mouse model, we identified transcriptomic indicators of SBS5 signature activation. Further studies are warranted to explore the connection between SBS5 and known oncogenic factors, such as tobacco smoking and PM2.5 exposure (Lau et al., 2022).

By integrating genomic and transcriptomic data from both human LUSC samples and a spontaneous LUSC mouse model, we established a direct association between SBS5 and central-type lung cancer. Moreover, we observed dynamic shifts in transcriptomic subtypes during LUSC progression, characterized by a transition from the secretory subtype to basal and classic subtypes. These findings provide preliminary evidence linking genomic mutations, transcriptomic profiles, and pathological subtypes. Future studies involving multi-omics analyses of patient samples will be essential to validate these observations in human LUSC.

Compared with LUAD, no specific targeted therapy has been developed for LUSC, underscoring the need to discover novel targets (Niu et al., 2022; Li et al., 2024). Herein, using our spontaneous LUSC mouse model, our study further revealed that late-stage disease progression was driven by specific genetic alterations, particularly mutations affecting the JNK signaling pathway (*e.g.*, *DACT1*) and cytoskeletal regulators (*e.g.*, *KIF26A*). Notably, previous studies revealed that JNK activity decreased during LUSC progression in both humans and mice, and *Jnk* knockout promoted LUSC progression (Liu et al., 2019), providing evidence that the JNK signaling pathway plays a critical suppressive role in LUSC progression. These findings provided potential targets for LUSC.

In summary, we delineated the genomic, transcriptomic, and cellular landscapes of LUSC progression, uncovering key molecular pathways and features. Our findings provided a valuable resource for studying LUSC development and identifying novel therapeutic targets. This research bridged the gap between tumor evolution, subtype classification, and therapeutic strategies, offering new insights for precision medicine in LUSC.

## Acknowledgements

This work was supported by grants to Dr. Jian Liu from the Natural Science Foundation (NSF) of China (General Grant: 82472637 and 82172899), the NSF of Zhejiang Province (Continuation Grant of Distinguished Young Scholars: LRG26H160001), Noncommunicable Chronic Diseases-National Science and Technology Major Project (2023ZD0502900/ 2023ZD0502902 and 2023ZD0507500/2023ZD0507501), Zhejiang University, the Open Fund of Zhejiang Provincial Key Laboratory of Pulmonology (KF202302), Zhejiang Provincial Key R&D Program of China (Grant No. 2025C02091), ZJE seed funding, ZJE 2024 International Campus Talent Special Funding Program, ZJE-UoE Joint Research Project, and Startup Funding of Tenure-track Assistant Professor of Zhejiang University. This work was also supported by Sanming Project of Medicine in Shenzhen (No. SZSM202403006) To Dr. Yong Tang. Principal Supervisor Jian Liu led the work. This work was in part supported by the Foundation from the Health Bureau of Zhejiang Province (2022KY804).

This work was supported by Zhejiang University-University of Edinburgh Institute (ZJE) and the Department of Respiratory and Critical Care Medicine, The Second Affiliated Hospital, Zhejiang University School of Medicine, Zhejiang University. We also acknowledge the help of members in JL’s lab. We thank the ZJE core facility, especially the ZJE mouse core facility, and the Biomed-X Laboratory of ZJE Institute, School of Medicine, Zhejiang University, for continuous support. We thanked Dr. Qifeng Li from Department of Pathology, Southern District People’s Hospital of Shenzhen to help the pathological evaluation of human and mouse samples. We also thanked Dr. Yongyong Shi and Dr. Zhiqiang Li from Department of Life Sciences and Technology, Shanghai Jiao Tong University to help revise our data and paper.

The Zhejiang University University of Edinburgh Institute (ZJE) undergraduate authors, such as Taoyu Zhu, Mingyang Xiao, Ran Ji, Huiran Wang, Xinyi Xu, Zhenting Zhang, and Bo Yang, gratefully acknowledged the profound impact of the ZJE HDRC3A (Human Disease: From Research to Clinic 3A) course, whose innovative pedagogical framework was instrumental in developing the critical perspective and interdisciplinary approach essential for this multi-omics work. The conceptualization and critical analysis presented in this review were directly shaped by the research-driven and omics-enhanced learning paradigm of the HDRC3A course. We would like to acknowledge the great help of undergraduate student Yixin Jiang in revising our paper. We also acknowledge the help of members in JL’s lab. We thank the Biomed-X Laboratory of Institute, School of Medicine, Zhejiang University, for continuous support.

The results shown here are in part based upon data generated by the TCGA Research Network: https://www.cancer.gov/tcga.

## Author contributions:CRediT

**Taoyu Zhu**: Data curation, Formal analysis, Investigation, Visualization, and Writing – original draft. **Yaping Xu**: Experiments, Validation, and Resources. **Junming Li**: Experiments, Validation, and Resources. **Zijin Wang**: Data curation, Formal analysis, and Investigation. **Mingyang Xiao**: Formal analysis, Investigation, and Visualization. **Boxin Liu**: Data curation, Formal analysis, and Investigation. **Boyu Wang**: Editing and Resources. **Manyu Xiao**: Editing and Resources. **Huiran Wang**: Visualization. **Xinyi Xu**: Visualization. **Zhenting Zhang**: Visualization. **Ran Ji**: Editing and Resources. **Bo Yang**: Visualization. **Songsong Li**: Editing and Resources. **Zuolin Shen**: Editing and Resources. **Xueqi Han**: Editing and Resources. **Xi Lu**: Editing and Resources. **Chen Lian**: Editing and Resources. **Xinyan Han**: Editing and Resources. **Yufei Liu**: Editing and Resources. **Silin Chen**: Editing and Resources. **Yunze Wang**: Editing and Resources. **Qian Tang**: Editing and Resources. **Yao Yao**: Editing and Resources. **Lian Wang**: Editing and Resources. **Huaqiong Huang**: Editing and Resources. **Qinglin Li**: Editing and Resources. **Da Wang**: Editing and Resources. **Xinwan Su**: Editing and Resources. **Yao Yao**: Editing and Resources. **Bing Xia**: Editing and Resources. **Hongshan Guo**: Supervision and Investigation, **Xushen Xiong**: Supervision and Investigation, **Xuru Jin:** Supervision, Writing – review & editing, and Funding acquisition. **Shirong Zhang**: Supervision, Writing – review & editing, and Funding acquisition. **Yong Tang**: Supervision, Writing – review & editing, and Funding acquisition. **Jian Liu**: Conceptualization, Funding acquisition, Resources, Supervision, and Writing – review & editing.

## Ethical statement

The Biomedical Research Ethics Committee of Zhejiang University approved all animal experiments (ZJU20240483). The Biomedical Research Ethics Committee of Shenzhen Nanshan People’s Hospital approved human samples (ky-2025-110602) and experiments.

## Data availability

The raw sequence data reported in this paper have been deposited in the Genome Sequence Archive in National Genomics Data Center, China National Center for Bioinformation / Beijing Institute of Genomics, Chinese Academy of Sciences (GSA: CRA031926, Reviewer link: https://ngdc.cncb.ac.cn/gsa/s/C6nDyxga; CRA023718, Reviewer link: https://ngdc.cncb.ac.cn/gsa/s/H5334z4u) that are publicly accessible athttps://ngdc.cncb.ac.cn/gsa. The published human scRNA-seq data were downloaded from Zenodo (https://doi.org/10.5281/zenodo.6411867) (Salcher et al., 2022).

## Code availability

Code relating to data processing and figure generation is available on Biocode (https://ngdc.cncb.ac.cn/biocode/) under Accession BT008025.

## Supplementary material

**Fig.S1 Basic information of TRACERX LUSC cohort patients, related to Fig.1**.

**A.** Patient timelines and mutational information in the TRACERX cohort. **B.** Bar plot showing the top 50 mutated genes in human TRACERX LUSC samples.

**Fig.S2 Alternative phylogenetic trees from stage I LUSC patients, related to Fig.1**.

**Fig.S3 Alternative phylogenetic trees from LUSC patients, related to Fig.1**.

**A.** Alternative phylogenetic trees from stage II LUSC patients. **B.** Alternative phylogenetic trees from stage III LUSC patients.

**Fig.S4 Validation of main findings in TCGA cohort LUSC patients, related to Fig.1**.

**A.** UMAP showed annotated major cell types in human LUSC samples. **B.** The Kaplan-Meier survival curves for overall survival (OS) in the TRACERX cohort lung cancer patients. The x-axis represented the time in months, while the y-axis showed the probability of survival. **C.** Bar plot showed the proportion of each cluster signature in distinct clusters. Error bars showed the 95% confidence interval for the calculated proportion.

**Fig.S5 DACT1 whole structure variation in human predicted by AlphaFold3, related to Fig.3**.

**Fig.S6 DACT1 whole structure variation in human predicted by AlphaFold3, related to Fig.3**.

**Fig.S7 NAIP whole protein structure variation in human and mouse predicted by AlphaFold3, related to Fig.3**.

**Fig.S8 KIF26A whole protein structure variation in human predicted by AlphaFold3, related to Fig.4**.

**A.** The box plot showing the expression level of *BDH1* in the TCGA database in pan-cancer. The dashed lines indicated the median values. A two-sided Wilcoxon test was applied for significance calculation. * p < 0.05, ** p < 0.01, *** p < 0.001. **B.** The Kaplan-Meier survival curves for OS in the TCGA cohort of LUSC patients. The x-axis represented the time in months, while the y-axis showed the probability of survival. **C.** KIF26A whole protein structure variation in human predicted by AlphaFold3

**Fig.S9 KIF26A protein structure variation in human and mouse predicted by AlphaFold3, related to Fig.4**.

**A.** KIF26A whole protein structure variation in human predicted by AlphaFold3. **B-C.** KIF26A whole protein structure variation in mouse predicted by AlphaFold3. **D-E.** KIF26A motor domain structure variation in human predicted by AlphaFold3.

**Fig.S10 KIF26A motor domain structure variation in human and mouse predicted by AlphaFold3, related to Fig.4**.

**A.** KIF26A motor domain structure variation in human predicted by AlphaFold3. **B-C.**

KIF26A motor domain structure variation in mouse predicted by AlphaFold3.

### Appendices

**Table 1 Conserved mutated genes and their mutation sites in human and mouse LUSC samples**

**Table 2 Impact of mutations on DACT1 and KIF26A total protein and domain structures**

**Table 3 Conserved DEGs between mouse and human genes and their variation levels**

**Table 4 Phase separation prediction score for KIF26A protein**

## Supplementary Tables

**Table S1 Detailed information of human chromosome coordinates, ref allele, alt allele, ref/alt allele counts for each called SNV and for coding SNV, as well as expected amino acid changes**

**Table S2 Detailed information of mouse chromosome coordinates, ref allele, alt allele, ref/alt allele counts for each called SNV and for coding SNV, as well as expected amino acid changes.**

**Table S3 The quality control data of human and mouse scRNA-seq data.**

