## Supplementary figures and images for "Polyclonal-Monoclonal Transition in Lung Squamous Cell Carcinoma Evolution"

### Supplementary Figure 1

Fig.S1. Basic information of TRACERX LUSC cohort patients, related to Figure 1.

A

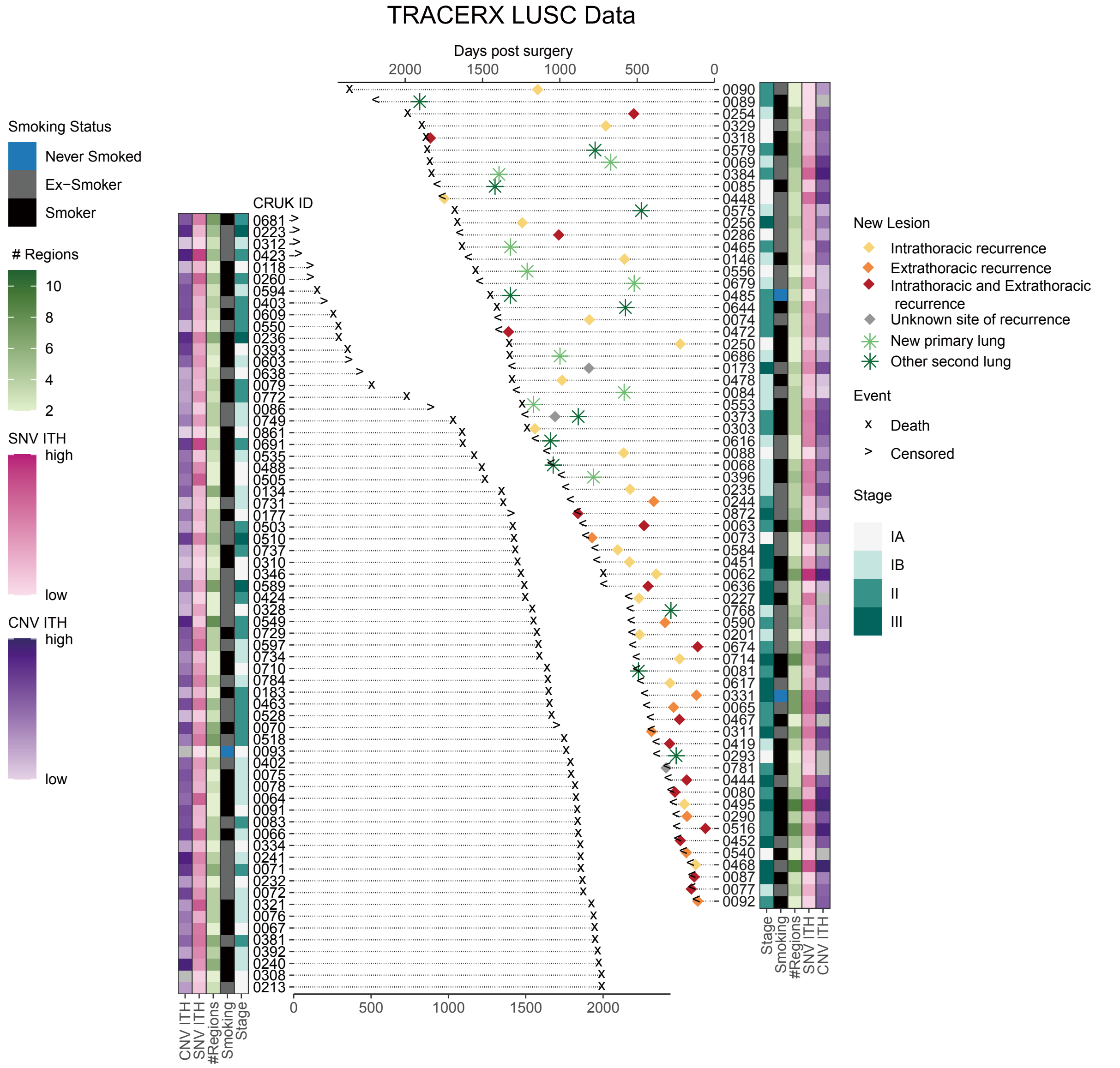


B

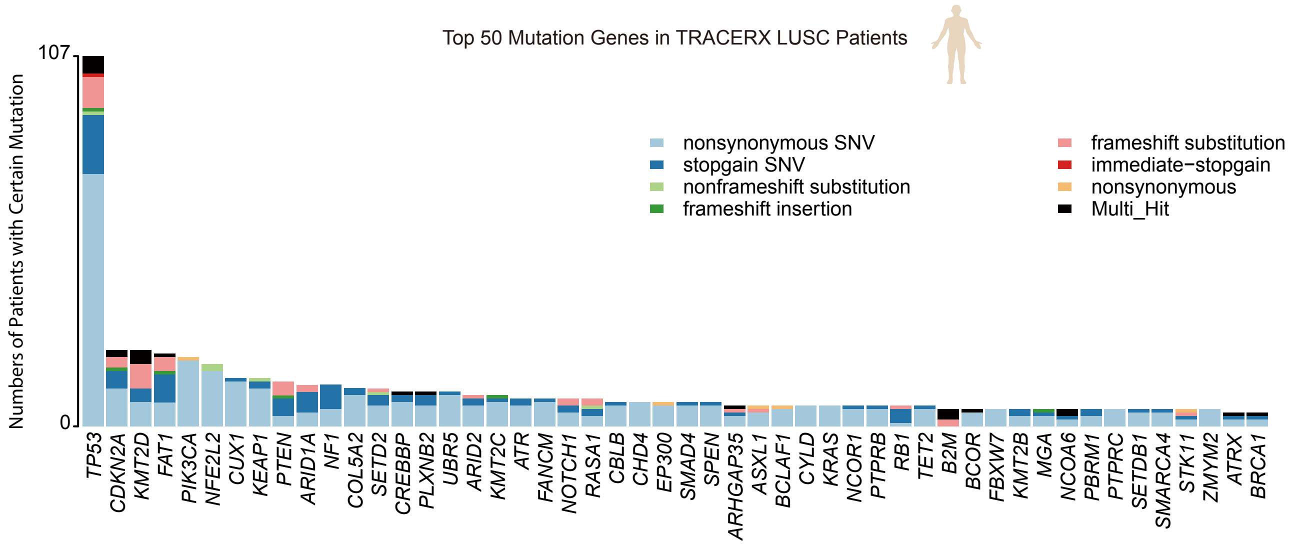

### Supplementary Figure 2

Fig.S2. Alternative phylogenic trees from stage I LUSC patients, related to Figure 1.

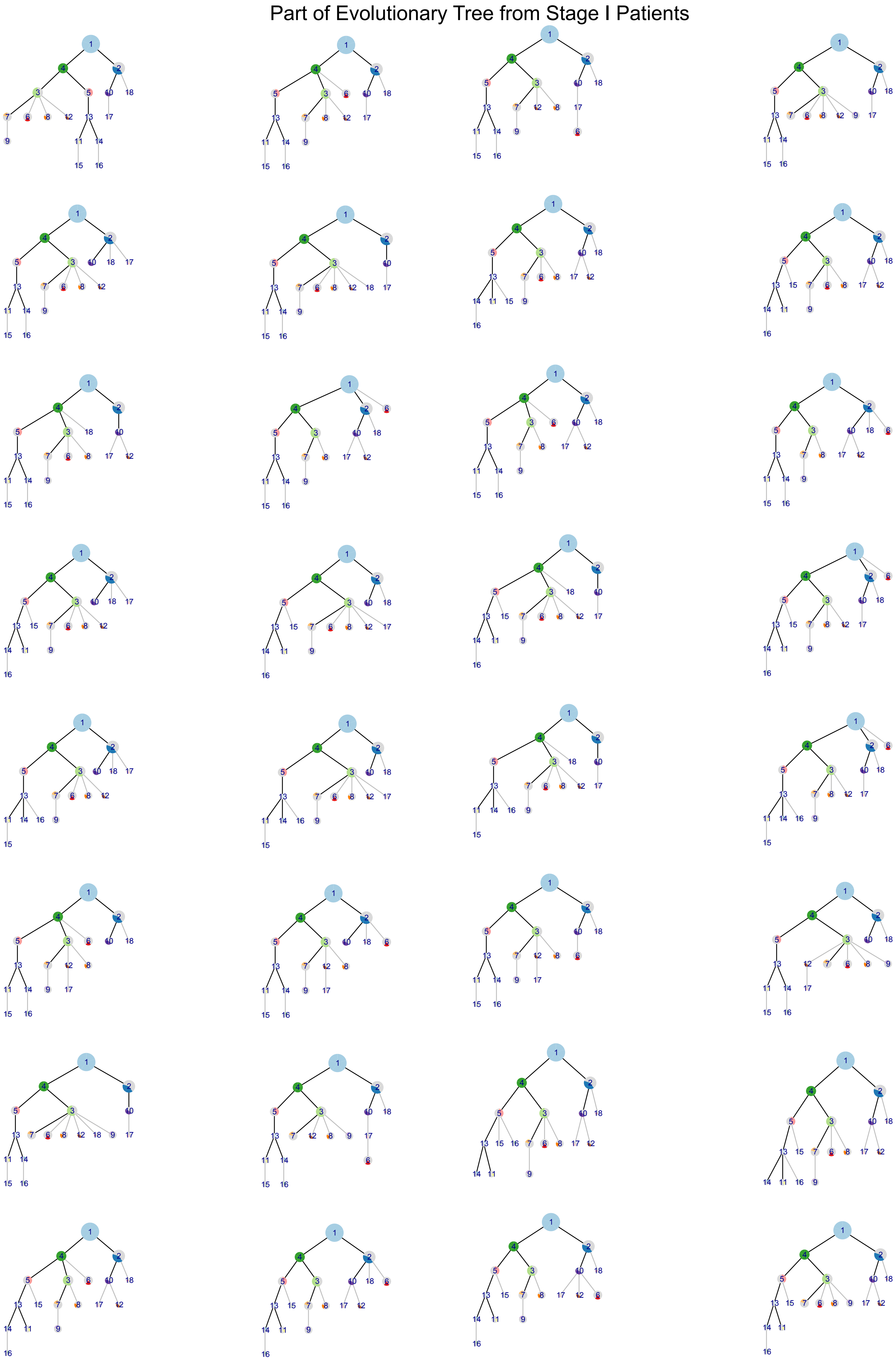

### Supplementary Figure 4

Fig.S4. Validation of main findings in TCGA cohort LUSC patients, related to Figure 1.

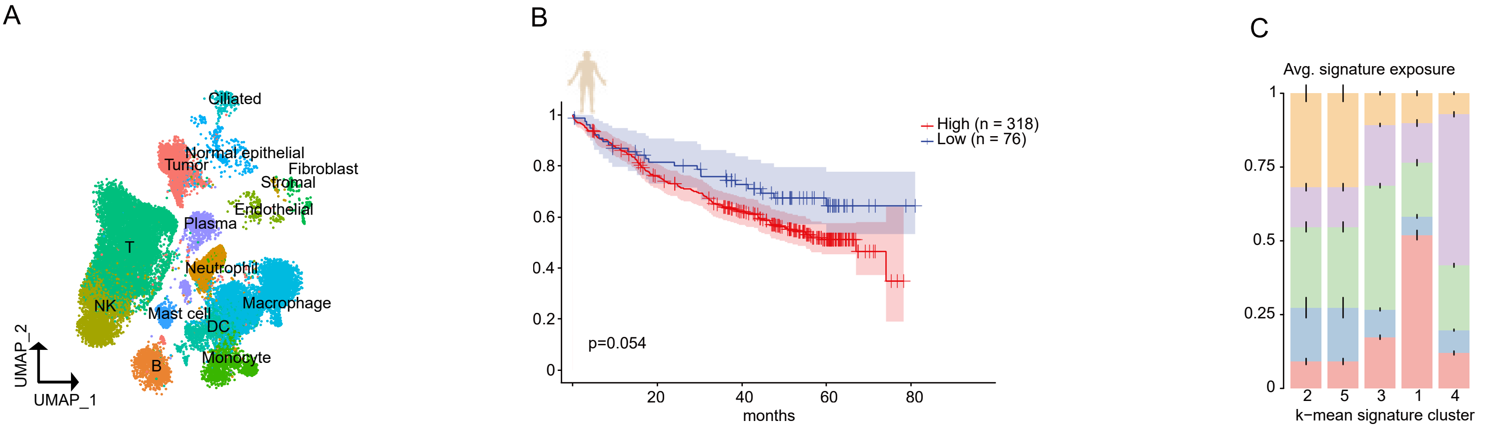

### Supplementary Figure 5

Fig.S5. Human DACT1 whole protein structure variation predicted by Alphafold3, related to Figure 3.

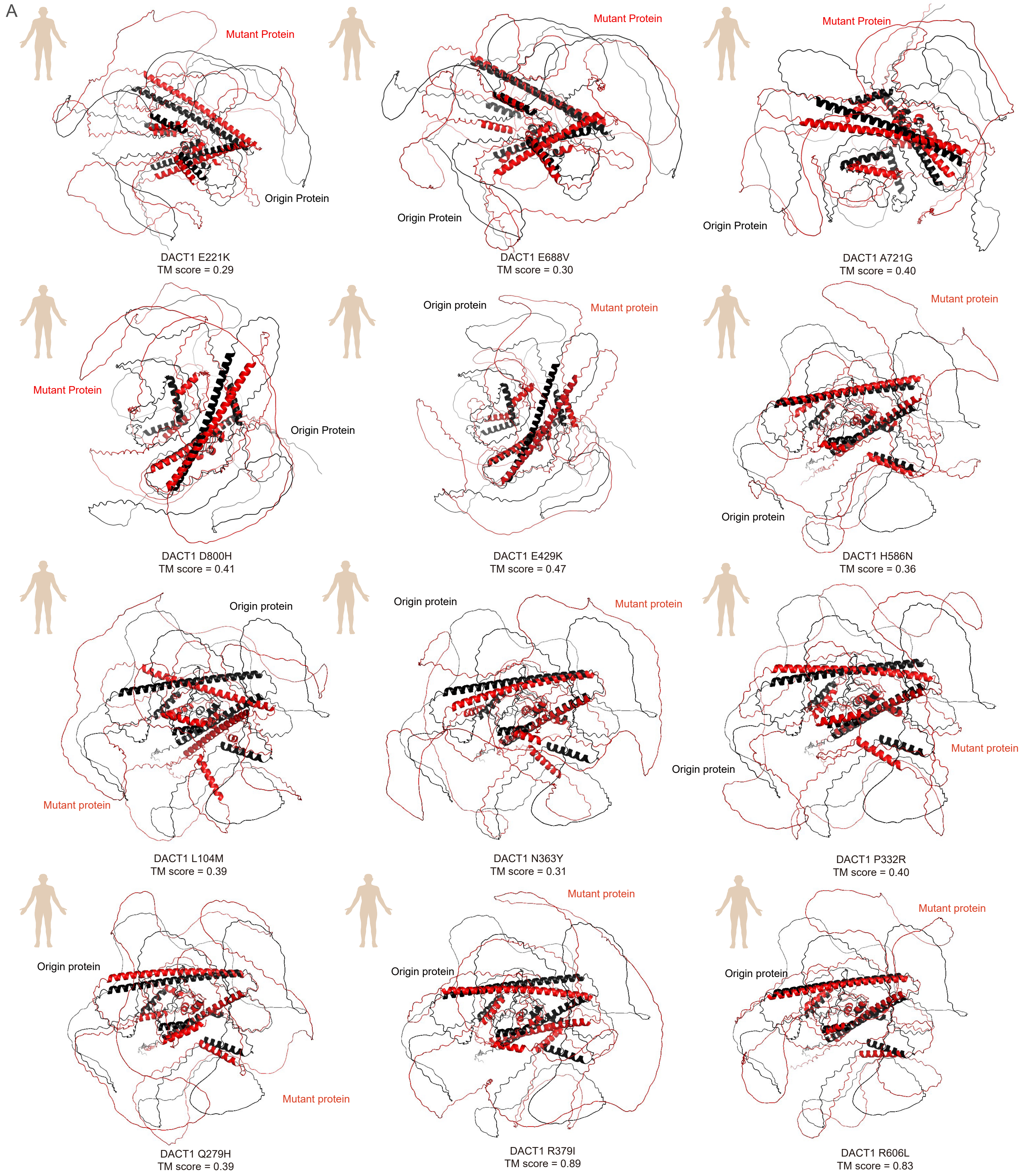

### Supplementary Figure 6

Fig.S6. Human DACT1 whole protein structure variation predicted by Alphafold3, related to Figure 3.

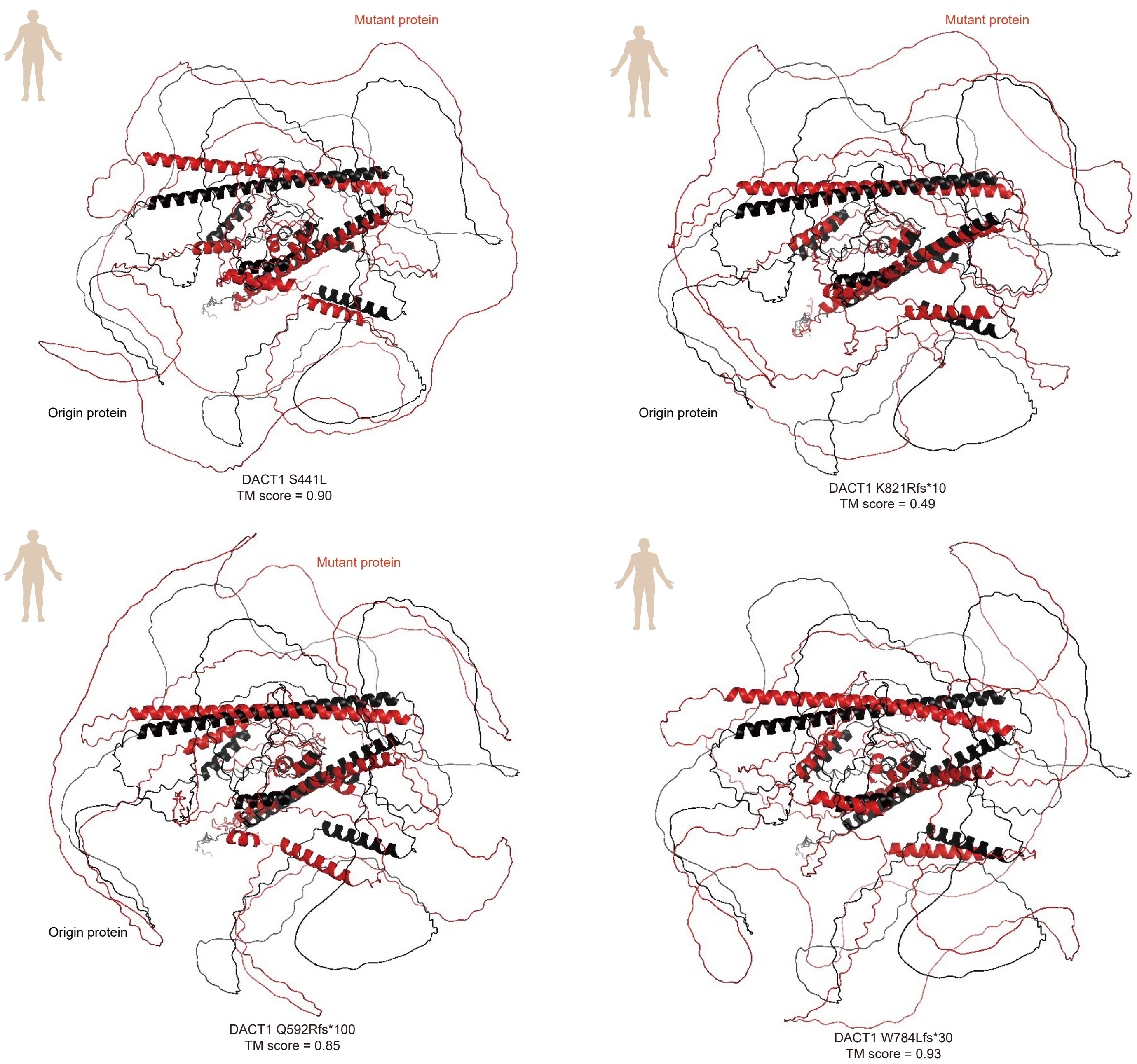

### Supplementary Figure 8

Fig.S8. Human KIF26A whole protein structure variation predicted by Alphafold3, related to Figure 4.

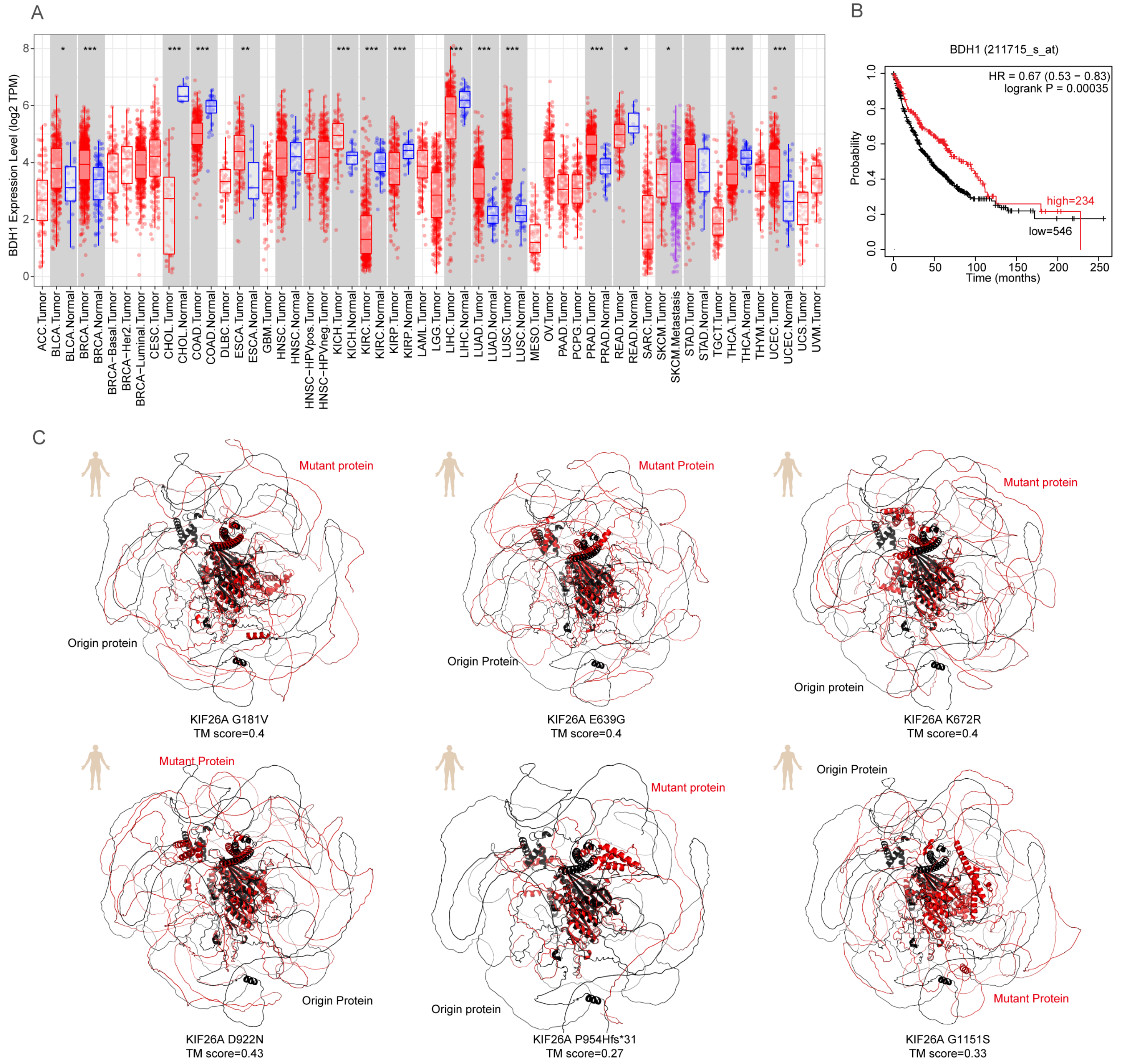
