## Supplementary Figure 3 for "Polyclonal-Monoclonal Transition in Lung Squamous Cell Carcinoma Evolution"

Fig.S3. Alternative phylogenic trees from stage I LUSC patients, related to Figure 1.

A

Part of Evolutionary Tree from Stage II Patients

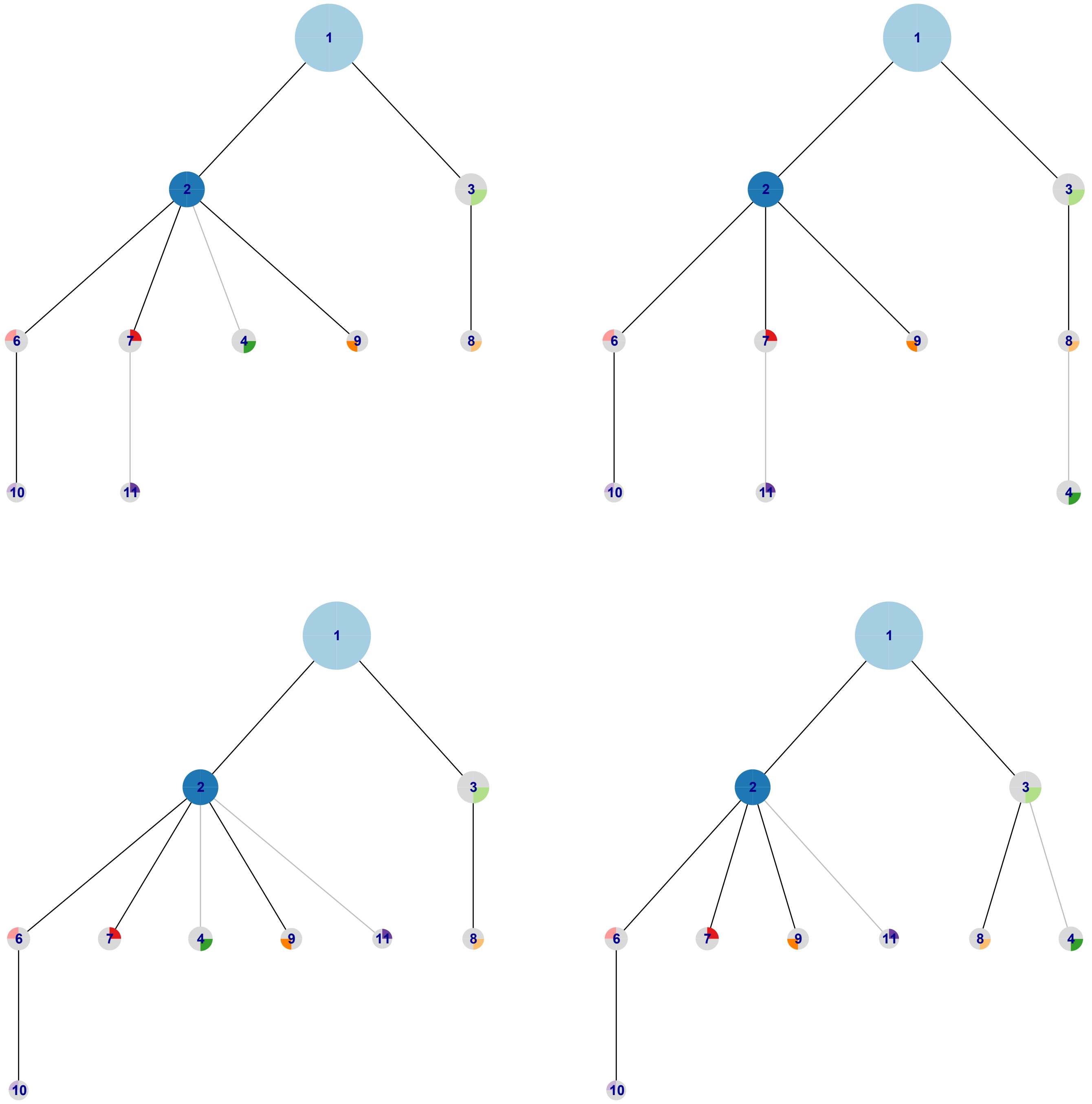


B

Part of Evolutionary Tree from Stage III Patients

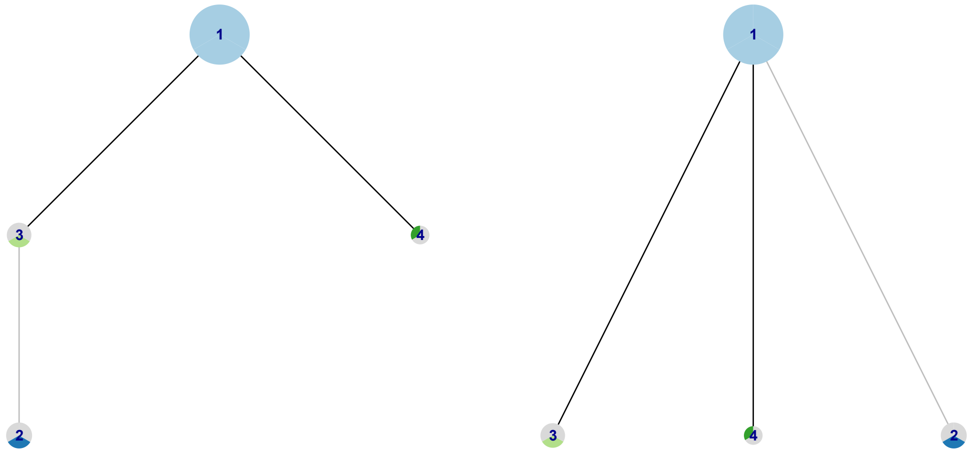
