## Supplementary Figure 7 for "Polyclonal-Monoclonal Transition in Lung Squamous Cell Carcinoma Evolution"

Fig.S7. Human NAIP whole protein and mouse NAIP1, NAIP2 whole protein structure variation predicted by Alphafold3, related to Figure 3.

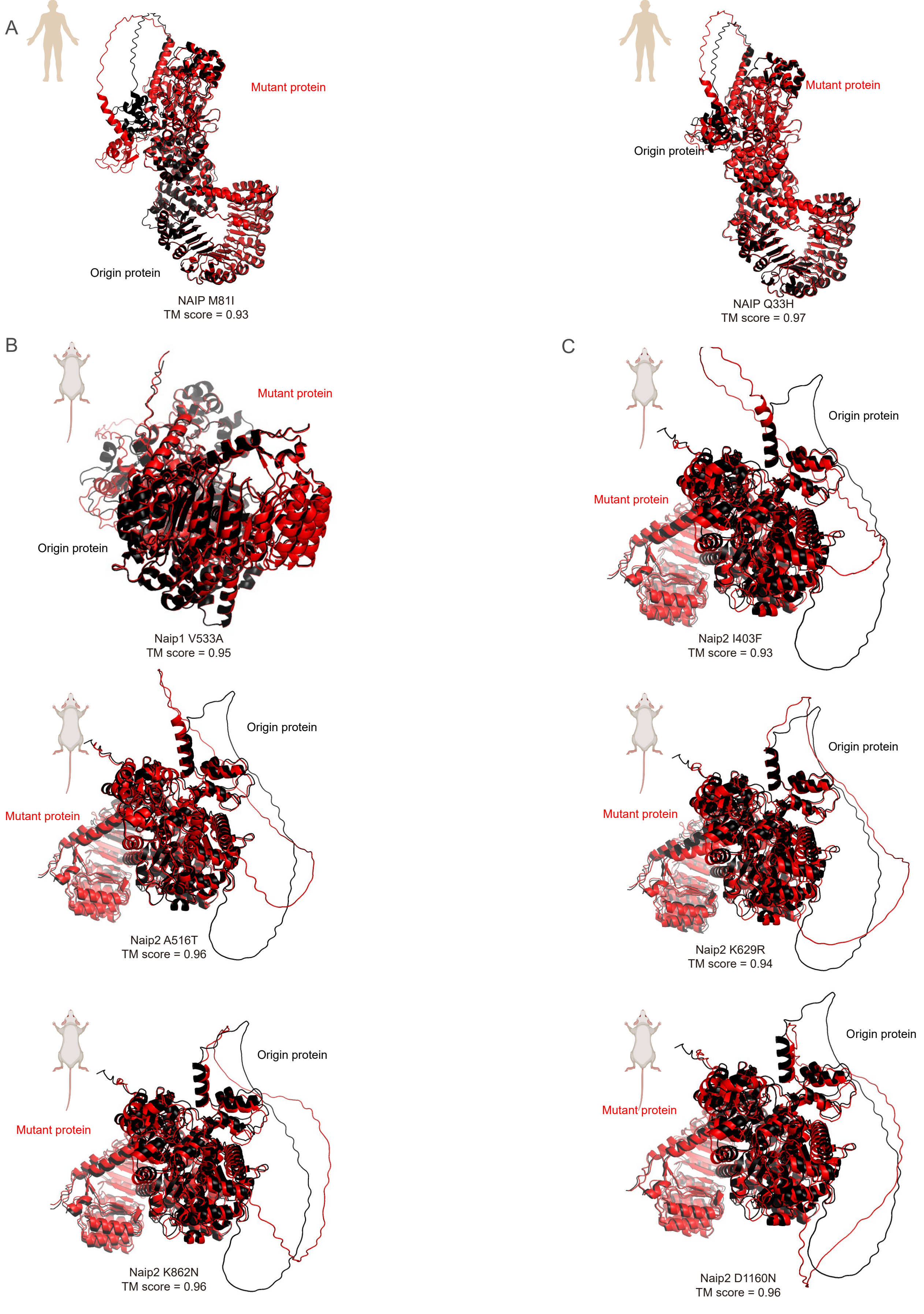
