## Supplementary Figure 9 for "Polyclonal-Monoclonal Transition in Lung Squamous Cell Carcinoma Evolution"

Fig.S9. Human and mouse KIF26A whole protein structure variation predicted by Alphafold3, related to Figure 4.

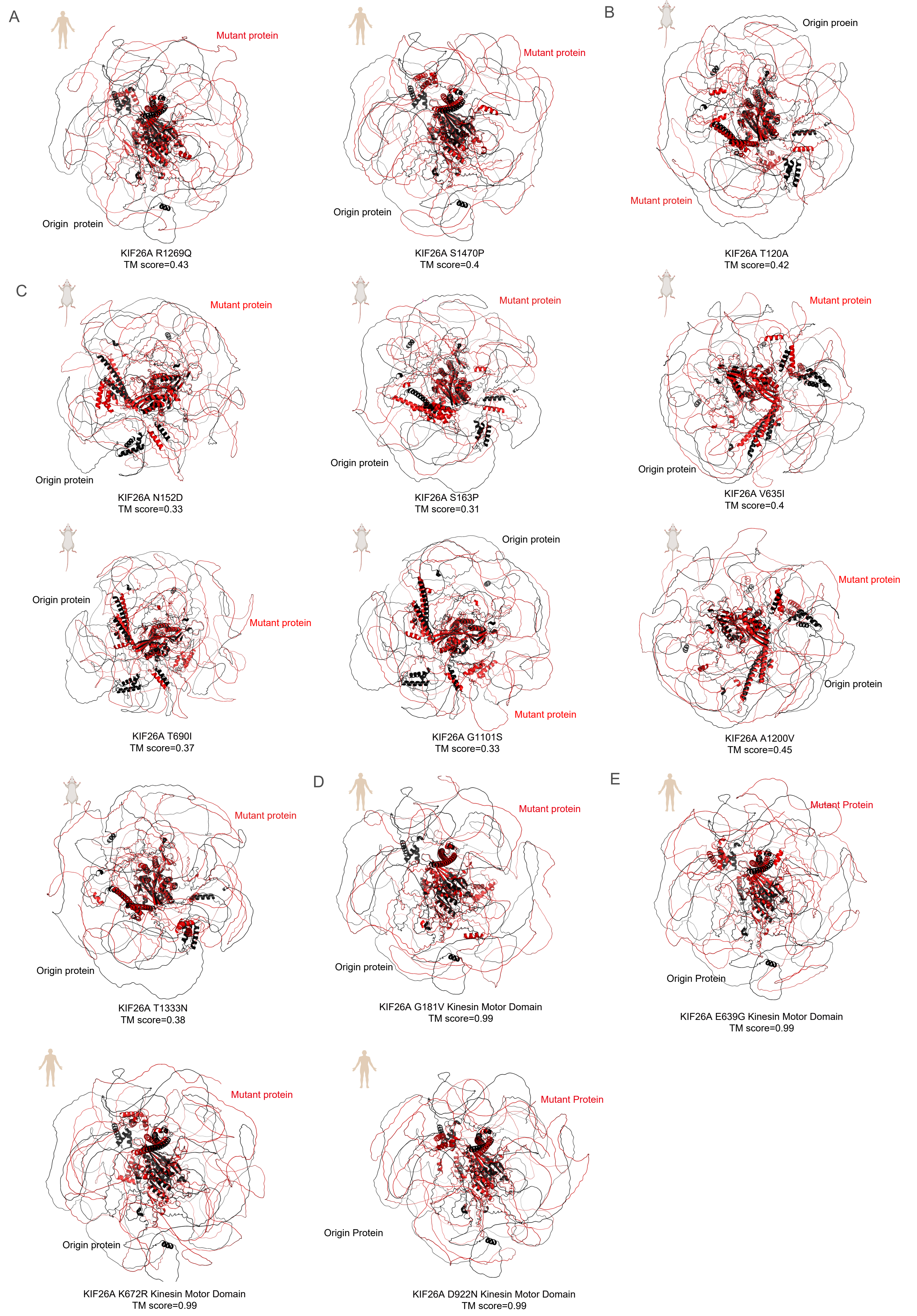
