## Supplementary Figure 10 for "Polyclonal-Monoclonal Transition in Lung Squamous Cell Carcinoma Evolution"

Fig.S10. Human and mouse KIF26A protein structure variation predicted by Alphafold3, related to Figure 4.

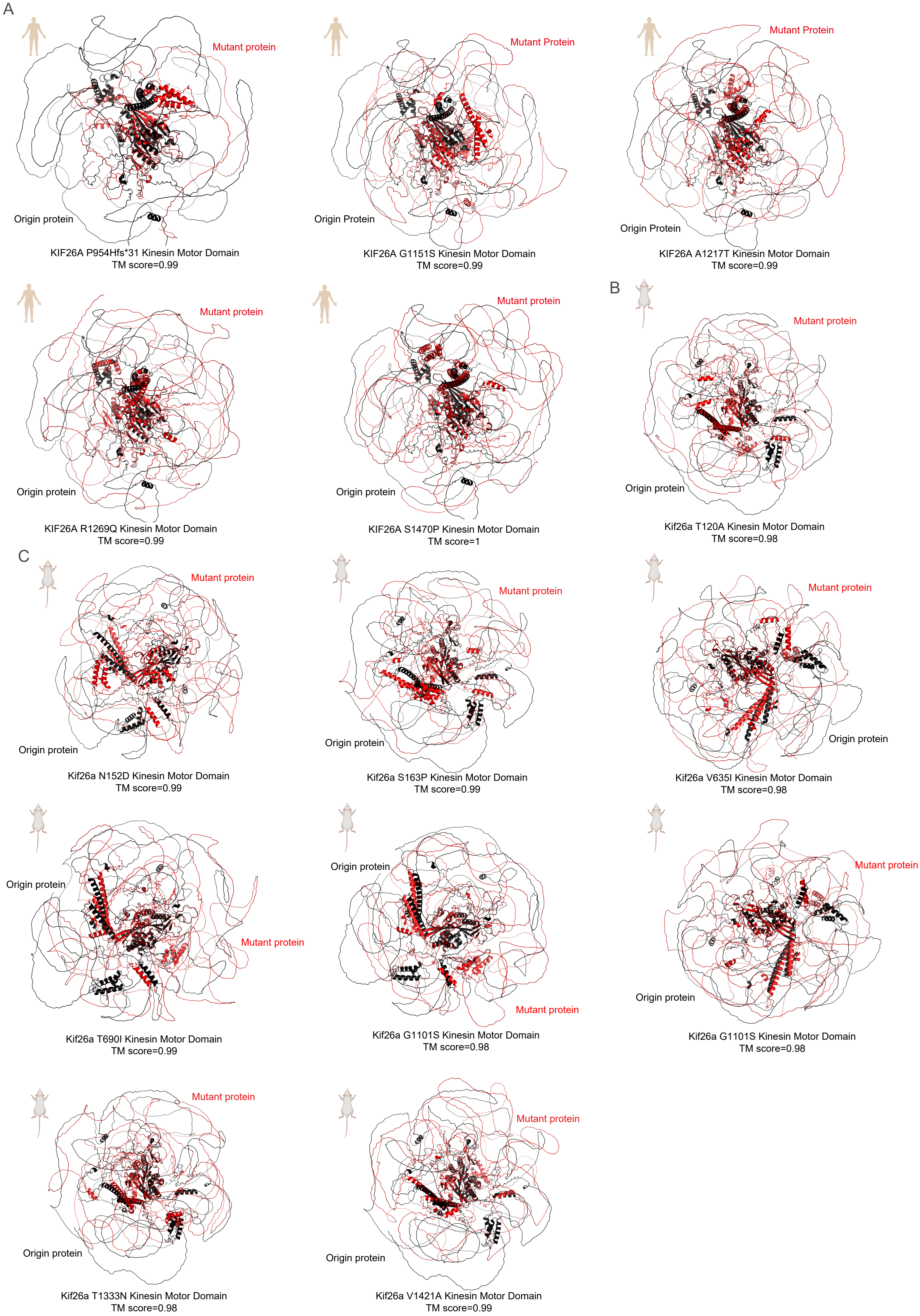
